# Developmental PBDE exposure impairs histamine release from mast cells by altering granule maturation and calcium signaling in adult male and female mice

**DOI:** 10.1101/2025.06.28.661534

**Authors:** Jared Franges, Lauren Malinowski, Chathuri De Alwis, Taylor Doolittle, Hannahlee Dixon, Yang Tang, Helen Watson, Jasmine Peace, Dereje Jima, Leslie Sombers, Gregory McCarthy, Heather Patisaul, Heather M. Stapleton, Natalia Duque-Wilckens

## Abstract

Polybrominated diphenyl ethers (PBDEs) are synthetic flame retardants once widely used in furniture, electronics, and other consumer products. Although phased out in the early 2000s, their persistence and recycling into new materials have led to continued environmental contamination and widespread human exposure —particularly through diet and indoor dust. Developing individuals face the highest exposure due to placental transfer, breastfeeding, and behavior, and are especially vulnerable to long-term effects. While developmental PBDE exposure has been linked to neurobehavioral, endocrine, and metabolic disruptions, effects on the immune system remain underexplored. To address this, we focused on mast cells—long-lived, tissue-resident innate immune cells enriched at barrier surfaces and perivascular sites throughout the body, including the brain. Their strategic positioning, broad receptor repertoire, and ability to rapidly release bioactive mediators suggest a key role in mediating multisystemic effects of developmental exposures. Here we show that maternal exposure to ∼87 μg/kg/day of PBDE throughout pregnancy and lactation—a dose aligned with the lower end known to affect metabolic and neurobehavioral outcomes in preclinical models—leads to persistent dysfunction in mast cell mediator release in adult male and female offspring. This was evidenced by blunted anaphylaxis-associated hypothermia and plasma histamine release *in vivo*. These deficits were not due to changes in tissue-resident mast cell numbers, but rather to an impaired capacity to sustain histamine release over time. In vitro studies of mast cells derived from adult bone marrow revealed that histamine synthesis was intact, but granule maturation and stimulus-induced calcium mobilization were disrupted, in association with downregulation of genes such as IGF2R, ITGA4, ITGB6, and NGFR. These results identify a novel mechanism by which developmental PBDE exposure impairs mast cell function, with implications for broader immune and physiological dysfunctions.

This is particularly concerning for developing individuals, who not only accumulate the highest levels via placental transfer, breastfeeding, and behavioral factors, but are also especially vulnerable to long-term effects. Despite well-documented impacts of developmental PBDE exposure on neurobehavioral, endocrine, and metabolic systems, the effects on the immune system remain comparatively underexplored. To begin addressing this gap, we focused on mast cells—innate immune cells well-positioned to contribute to the multisystemic effects of developmental exposures. Mast cells are long-lived, tissue-resident cells enriched at barrier surfaces and perivascular sites throughout the body, including the brain. Their widespread distribution, extensive receptor repertoire, and unique ability to store and rapidly release bioactive mediators from cytoplasmic granules position them as key modulators of immune, endocrine, and nervous system function. Using oral exposure to two doses of a PBDE mixture throughout pregnancy and lactation in mice, here we show that maternal exposure to ∼87 μg/kg/day—aligned with the lower end of doses known to affect metabolic and neurobehavioral outcomes in preclinical models, and within 10-fold of levels measured in human serum and placenta—leads to persistent dysfunction in mast cell mediator release in adult male and female offspring. This was evidenced by blunted anaphylaxis-associated hypothermia and plasma histamine release *in vivo*. These deficits were not due to changes in tissue-resident mast cell numbers, but rather to an impaired capacity to sustain histamine release over time. Studies in bone marrow–derived mast cells (BMMCs) revealed that histamine synthesis was intact, but granule maturation and stimulus-induced calcium mobilization were disrupted, in association with downregulation of genes such as IGF2R, ITGA4, ITGB6, and NGFR. Given that the bone marrow is the primary postnatal source of mast cells, these findings suggest that PBDEs induce lasting reprogramming at the level of hematopoietic progenitors—with broad implications not only for mast cell function across tissues, but potentially for other immune cell lineages as well. In sum, this study provides the first evidence that developmental exposure to PBDEs induces long-lasting impairments in mast cell functions, suggesting a previously unrecognized mechanism by which early-life exposure to environmental toxicants could contribute to persistent physiological and behavioral dysfunctions

## Introduction

Polybrominated diphenyl ethers (PBDEs) are a class of synthetic flame retardants that contain a diphenyl ether backbone with one to 10 substituted bromine atoms around the aromatic moiety. Up until the early 2000s, three commercial mixtures of PBDEs, known as PentaBDE, OctaBDE and DecaBDE^1^, were manufactured and commonly used in furnishings, electronics and some consumer goods to meet specific fire safety standards. Examples include upholstered furniture, televisions and building materials. Although restrictions on the use of PBDEs began in the early 2000s due to their persistence and toxicity^2,3^, they continue to pose a significant public health and environmental concern. Many discarded products containing PBDEs are deposited in landfills, where PBDEs gradually leach out of the product matrix into surrounding soil and water due to their lack of chemical binding to the material substrate^4,5^. Combined with their resistance to degradation and strong lipophilicity, this leaching contributes to widespread bioaccumulation across ecosystems and biomagnification through the food chain^6–12^. Additionally, while the production of PBDE-containing products has declined, many continue to be recycled into new materials—resulting in PBDEs being inadvertently carried over into consumer items, including children’s toys^13–15^, kitchen utensils and food packaging materials^16^ As a result, despite regulatory restrictions, human exposure remains widespread—primarily through diet and through the ingestion and inhalation of contaminated house dust^17,18^—and PBDEs continue to be detected in human tissues^19–23^.

This is particularly concerning for infants and toddlers, who are not only at greater risk of adverse health effects from PBDE exposure due to ongoing tissue development, but also tend to accumulate several-fold higher serum PBDE concentrations as compared to adults^24–26^. This increased PBDE burden is driven by placental and lactational transfer^23,27–32^, as well as child-specific behaviors such as frequent hand-to-mouth activity and close contact with dust-contaminated surfaces^20,33^. Developmental PBDE exposure has been linked to a broad spectrum of adverse outcomes in both human and animal studies—including endocrine disruption^34–38^, reproductive deficits^35,39^, metabolic disturbances^40–43^, and cognitive and social impairments^44–46^—suggesting that these chemicals target core regulatory systems with widespread physiological impact.

Mast cells are well-positioned to contribute to these multisystem effects. These long-lived, tissue-resident immune cells are distributed throughout the body-including the brain-and are particularly enriched at barrier surfaces and perivascular sites. Their strategic localization, coupled with an extensive receptor repertoire, enables mast cells to sense and respond to diverse physiological signals, including cytokines and other immune mediators^47–51^, hormones^52–56^, neurotransmitters^57–59^, and environmental cues^60–63^. This responsiveness is amplified by their unique ability to store a variety of bioactive mediators in cytoplasmic secretory granules. These include proteases such as tryptase and chymases, heparin, tumor necrosis factor (TNF), vascular endothelial growth factor (VEGF), and notably, amines such as serotonin and histamine —the most extensively studied and one of the most abundant small-molecule mediators stored in mast cells^64,64–71^.

Histamine is a biogenic amine consisting of an imidazole ring attached to an ethylamine chain. Although it can be synthesized by various cell types, including neurons, macrophages, and platelets, mast cells—in conjunction with basophils to a lesser extent—are the primary source of preformed histamine in the body^67,72^. Mast cells uniquely possess the enzymatic machinery and specialized granules required to store histamine– alongside other pre-stored mediators– at high concentrations. This enables them to release histamine not only rapidly, but also in a sustained manner, positioning them as key responders in diverse physiological and pathophysiological contexts. Once released, histamine exerts pleiotropic effects across tissues by acting through four G protein–coupled receptors (H1R–H4R). For example, histamine regulates vascular tone and permeability^73–75^, shapes immune cell recruitment and activation^76–78^, and helps protect immune cells against oxidative stress–induced damage^79^. It also influences gonadal function and hormone secretion^80–83^, supports liver ketogenesis^84^, and contributes to endothelial barrier dysfunction^85^. In the brain—where mast cells account for approximately 50% of total histamine^86,87^— mast cell–derived histamine has been implicated in promoting arousal^88^, modulating baseline anxiety^87,88^, contributing to microglial activation^75,89^, organization of sex-specific neural circuits^90^, and modulating the stress response via interactions with corticotropin-releasing factor (CRF) neurons^91,92^.

Given histamine’s broad physiological roles across nervous, immune, and endocrine systems —and the fact that mast cells serve as a primary systemic reservoir—disruptions in mast cell histaminergic activity can have widespread consequences. We hypothesized that developmental exposure to a PBDE mixture would induce long-lasting dysregulation of mast cell histamine release. To test this, we administered daily oral doses of a PBDE mixture—composed of predominant congeners from commercial penta-and deca-BDE formulations—at either 9.97 μg/kg (“low” dose) or ∼87 μg/kg (“high” dose) to mouse dams throughout mating, gestation, and lactation. Mast cell physiology was then assessed in the adult offspring. High-dose PBDE exposure led to persistent impairments in histamine release following both IgE-dependent and IgE-independent stimulation in male and female offspring. These deficits were not due to altered histamine synthesis or storage, but rather to impaired granule maturation and sustained calcium-dependent exocytosis. Notably, these alterations were evident in both tissue-resident mast cells and bone marrow–derived mast cells (BMMCs), indicating that PBDEs reprogram mast cell progenitors during development and disrupt function across tissues into adulthood.

## Methods

### Animals

All animal procedures were conducted in accordance with the U.S. Animal Welfare Act and the DHHS Guide for the Care and Use of Laboratory Animals, and approved by the Institutional Animal Care and Use Committee (IACUC) at North Carolina State University (NCSU). Procedures were supervised by a university veterinarian. Breeding pairs of C57BL/6J mice (Jax Strain #000664) were used to establish an in-house colony. Mice were housed in a humidity-and temperature-controlled facility with 12-h:12-h light:dark cycles (25 °C; 45–60% humidity), and provided ad libitum access to glass-bottled water and a soy-free diet (2020X Teklad Global Soy Protein-Free Extruded Rodent Diet). Housing included woodchip bedding (Beta chip), enrichment materials (cinkl-nest, nestlets, tunnels from Bio-Serv), and thoroughly washed polysulfone caging. Pups were weaned at postnatal day (PN) 21, ear-tagged, and housed in same-sex groups of 2–4 animals.

### Dosing

Eight-week-old females were paired with males and dosed daily from the first day of pairing through PN21. The dosing solution consisted of vehicle (sesame oil) or one of two PBDE mixtures, formulated by Dr. Heather Stapleton’s lab^30^, containing BDE-28,-47,-99,-100,-153, and-209 **(Fig. 1A).** These congeners represent the most abundant components of commercial penta-and deca-BDE formulations and were prepared to mimic the average distribution in US house dust. Neat PBDEs were dissolved in pure sesame oil to prepare solutions and concentrations were confirmed by gas chromatography mass spectrometry. Dosing began at the time of mating and continued throughout gestation and lactation. Each day, 10 μL of the assigned solution (vehicle control, low PBDE, or high PBDE) was pipetted onto mini marshmallow bits (Jet-puffed), which the dams readily consumed (**Fig. 1B**). The dosing was administered by an experimenter who was aware of the solution labels (A, B, and C) but blinded to their content. The low dose (∼9.97 μg/kg) was selected to approximate, and likely fall within up to 10-fold of, PBDE levels measured in human serum and placenta^93,94^. The high dose (∼87 μg/kg) was based on prior rat studies demonstrating placental accumulation at levels ∼10–100-fold higher than typical human exposures^30^—a range commonly used to model the safety margin built into regulatory risk —and also aligns with the lower end of doses used in previous mouse studies assessing metabolic and neurobehavioral outcomes of developmental PBDE exposure^95–97^. Pups were weaned at postnatal day (PN) 21 and left undisturbed until adulthood (PN56). Each litter (n = 5 per treatment group, average 7 pups per litter) was divided across four different experiments (behavior test, passive systemic anaphylaxis, peritoneal mast cell isolation, bone marrow isolation), with no more than two animals per sex per litter included in any one experiment to account for potential litter effects.

**Figure 1.**
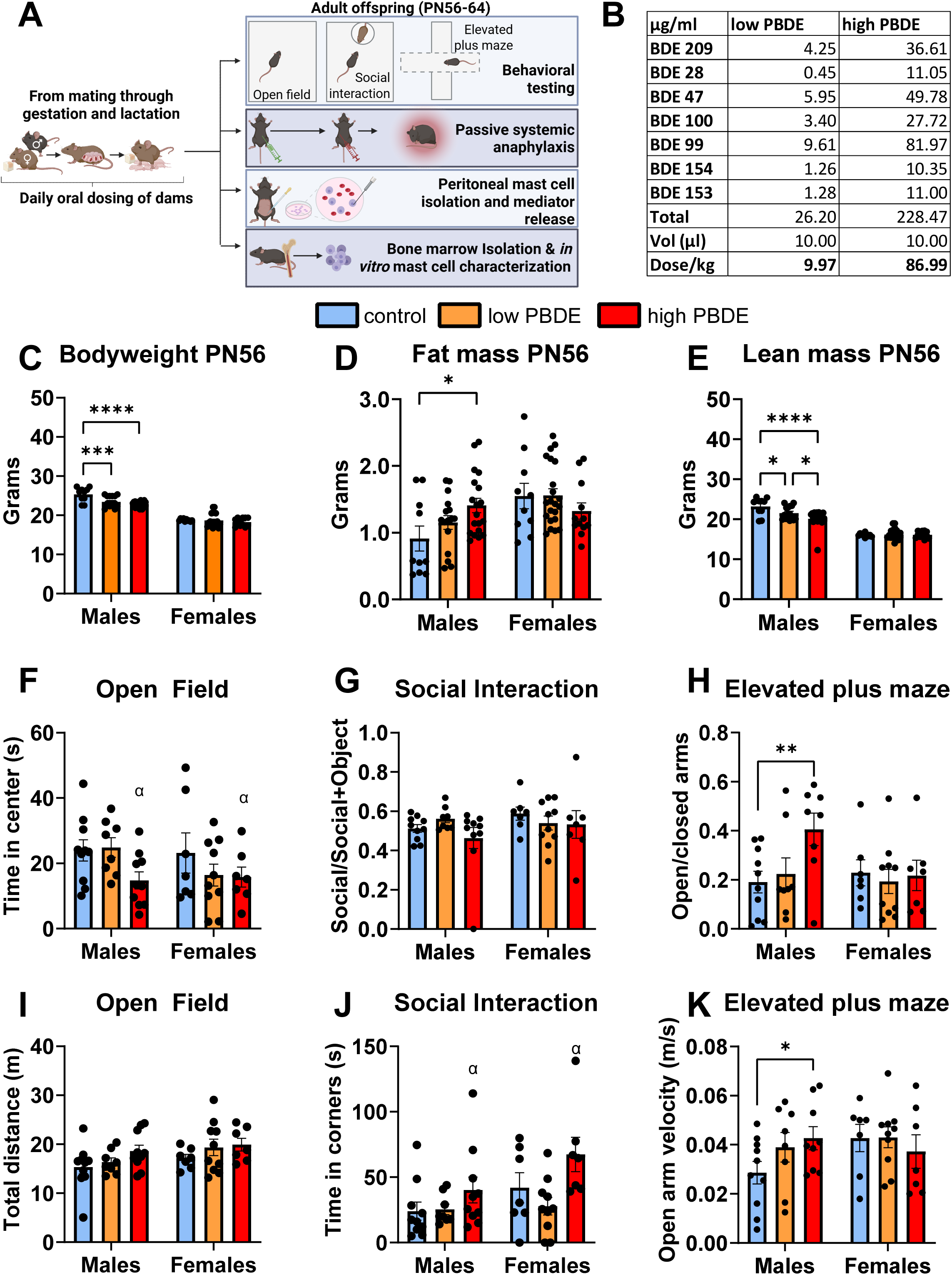
Developmental PBDE exposure alters body composition and behavior in a sex-specific manner. **(A)** Experimental timeline. **(B)** Composition of the low and high PBDE dosing mixtures, including individual congener concentrations and total administered dose. **(C–E)** Body composition at PN56 (n = 10–12/group). Two-way ANOVA revealed significant main effects of sex (F (1, 84) = 354.9, p < 0.0001), treatment (F (2, 84) = 8.233, p = 0.0005), and a sex × treatment interaction (F (2, 84) = 4.025, p = 0.02) on bodyweight **(C).** Fisher’s LSD showed that this effect was specific to males, with both low (p < 0.0004) and high (p < 0.0001) PBDE groups weighing significantly less than controls. This reduction in bodyweight was accompanied by male-specific changes in body composition. Fat mass **(D)** showed a main effect of sex (F (1, 83) = 8.935, p = 0.004) and a sex × treatment interaction (F (2, 83) = 3.753, p = 0.03), driven by a significant increase in fat in high PBDE–exposed males compared to controls (Fisher p = 0.01). Lean mass **(E)** showed main effects of sex (F (1, 83) = 249.8, p < 0.0001), treatment (F (2, 83) = 5.618; p = 0.005), and sex × treatment interaction (F (2, 83) = 5.804, p = 0.004), driven by significant reductions in lean mass in both low (p = 0.01,) and high (p < 0.0001) PBDE males relative to controls. **(F–K)** Behavior at PN56 (n = 7–10/group). **Open field test**: A trend for a main effect of treatment was observed for time spent in the center (**F**; F(2,48) = 2.8, p = 0.07), although Fisher LSD revealed reduced center time in high PBDE vs control mice (p = 0.02), independent of sex. No treatment effects were detected for total distance traveled in the open field **(I). Social interaction test**: No sex or treatment effects were observed on the social/social+object ratio **(G).** However, there was a significant main effect of treatment (F(2,46) = 5.08, p = 0.01), and sex (F(1,46) = 4.7, p = 0.03) on time spent in corners **(J)**, driven by increased corner time in high PBDE vs control mice (p = 0.03), independent of sex.**Elevated plus maze**: No sex, treatment or interaction effects were detected in the open arms/total time **(H).** Nonetheless, Fisher LSD indicated that high PBDE males spent less time in the open arms compared to controls (p = 0.008). This was accompanied by increased velocity in the open arms in high PBDE males (Fisher LSD p = 0.047, no significant effects detected in two-way ANOVAS) compared to controls **(K).**

### Body composition

Body composition was evaluated in a randomly selected subset of adult offspring (PN52–PN55) from each treatment group using EchoMRI™-100H Body Composition Analyzer (Echo Medical Systems, Houston, TX), which provides non-invasive measurements of lean mass, fat mass, free water, and total body water in live, unanesthetized mice. Prior to scanning, mice were gently restrained in an appropriately sized clear plastic tube provided by the manufacturer to minimize stress and ensure proper positioning. Each scan lasted approximately 1–2 minutes per animal. To avoid potential confounding effects of body composition on downstream analyses, the animals assessed for body composition were evenly distributed across the experimental endpoints. All assessments were conducted during the light phase, and experimenters were blinded to exposure group during data acquisition and analysis.

### Behavior Testing

Behavioral assessments were conducted in a dedicated behavioral suite under red light conditions between 1:00 PM and 5:00 PM to minimize circadian influences on activity. Testing occurred over two consecutive days in the following order: open field test (day 1), social interaction test (after open field, day 1), and elevated plus maze (day 2). Prior to testing each day, mice were habituated to the testing room for at least 60 minutes. All behavior was recorded and automatically analyzed using ANY-maze™ software (Stoelting Co., Wood Dale, IL). All apparatuses were thoroughly sanitized with 70% ethanol before the start of testing and between subjects to eliminate olfactory cues and ensure consistency across trials. <u>Open Field Test:</u> The open field test was used to assess general locomotor activity and anxiety-like behavior^98^. Mice were individually placed in the center of a white Plexiglas arena (90 cm × 45 cm × 45 cm) and allowed to explore freely for 3 minutes. The arena was divided into center and peripheral zones using ANY-maze software. Total distance traveled, time spent in the center, and number of entries into the center zone were used as indices of locomotion and anxiety-like behavior. <u>Social Interaction Test:</u> The social interaction test is a widely used behavioral assay to assess an animal’s motivation to engage with a novel social stimulus versus a non-social object^99,100^. Immediately following the open field test—which also served as acclimation to the testing environment—mice were briefly returned to their home cage while the arena was sanitized with 70% ethanol. A metal mesh cage enclosure (9 cm diameter × 18 cm height) was then placed in one corner of the open field arena to serve as the stimulus cage. The test consisted of two consecutive 3-minute phases: (1) Acclimation Phase – the empty mesh cage was present and the subject mouse was reintroduced into the arena to assess baseline exploratory behavior; (2) Interaction Phase – an unfamiliar, age-and sex-matched C57BL/6J stimulus mouse was placed into the enclosure, and the subject mouse was allowed to interact freely. Time spent in the defined interaction zone (a 2 cm perimeter around the enclosure) was automatically recorded. The ratio of time spent in the interaction zone during interaction divided by the time in interaction zone during acclimation and interaction was used as an index to measure social motivation. <u>Elevated Plus Maze:</u> The elevated plus maze was used to assess exploratory and anxiety-like behavior^101^. The apparatus consisted of two open arms and two closed arms (35 cm × 5 cm × 15 cm) arranged in a plus-shape and elevated 50 cm above the floor. Mice were placed in the center of the maze facing an open arm and allowed to explore for 5 minutes. The number of entries, velocity, distance, and total time spent in open vs. closed arms were automatically recorded by Anymaze. Anxiety-like behavior was indexed as the ratio of time spent in open versus closed arms, with lower ratios interpreted as increased anxiety-like behavior.

### Passive systemic anaphylaxis

mice were sensitized with an intraperitoneal (IP) injection of 5 μg of anti-DNP IgE (SPE-7, Sigma-Aldrich) in 100 μL of sterile saline. 24h later, the mice were challenged with an IP injection of 50 μg of DNP-HSA (Sigma-Aldrich) in 100 μL of saline, as previously described^102,103^. Rectal temperatures were recorded right before DNP-HAS injection and every 5 minutes for up to 30 minutes following the DNP challenge using a TH-5 Thermalert thermometer (Physitemp, Clifton, NJ), after which animals were immediately euthanized for tissue collection.

### Plasma histamine

Circulating histamine levels were measured from plasma collected at euthanasia using a competitive enzyme-linked immunosorbent assay (ELISA) kit (EA31; Oxford Biomedical Research), following the manufacturer’s protocol. All samples were run in duplicate. Plates were read on a microplate reader at the recommended wavelength, and histamine concentrations were calculated using a standard curve generated in parallel.

### Assessment of tissue mast cell numbers and activation status

To evaluate tissue-resident mast cells, small intestinal mesentery windows and dura mater were dissected, whole-mounted on glass slides, fixed in Carnoy’s fixative (ethanol:chloroform:acetic acid, 6:3:1), and stained with 0.1% toluidine blue (Sigma-Aldrich) for 30 minutes, as previously described^104^. Excess stain was rinsed with distilled water and slides were air-dried. Mast cells were identified by their metachromatic granules and counted in four non-overlapping fields per tissue at 10× magnification. Each field encompassed the full high-power field of view. Quantification was performed by a blinded observer using ImageJ software. At least four mesenteric windows and dura regions per mouse were analyzed to ensure reproducibility and minimize sampling bias.

### Fast-scan cyclic voltammetry for quantitative co-detection of histamine and serotonin

Peritoneal mast cells were isolated from adult male and female offspring perinatally exposed to either vehicle control or high-dose PBDEs. Immediately following sacrifice, 5–10 mL of pre-warmed buffer (150 mM NaCl, 5 mM KCl, 1.2 mM MgCl_2_, 5 mM glucose, 10 mM HEPES, and 2 mM CaCl_2_ at pH 7.4) was injected into the peritoneal cavity. After gently massaging the abdomen for 1–2 minutes to dislodge resident cells, the cavity was carefully opened with a sterile scalpel, and the lavage fluid containing peritoneal cells was collected into sterile centrifuge tubes. Samples were centrifuged at 600 ×Lg for 5 minutes at 37 °C to pellet the cells, which were then resuspended in complete mast cell culture media and plated into culture dishes. Dishes were incubated at 37L°C for at least 3 hours to allow cell adherence prior to experimentation. For exocytosis recordings, the media was replaced with ∼3 mL of pre-warmed buffer, and all recordings were conducted at 37 °C. A 34-μm diameter disk carbon-fiber microelectrode was gently placed in contact with the mast cell membrane to directly monitor exocytosis, and a Ag/AgCl reference electrode was placed in the far side of the dish. The potential was swept from +0.2 V to +1.3 V (1000 V/s) before a negative sweep to-0.1 V followed by another positive sweep to +0.2 V where the potential was held constant for ∼97 msec. This waveform was repeated at a frequency of 10 Hz. Vesicular histamine and serotonin release was evoked by focal application of compound 48/80 (Sigma) using a picospritzer. This voltammetric method allows for high-resolution, real-time identification and quantification of histamine and serotonin release events from single mast cells. Peaks evident in the current vs time traces with intensity exceeding five times the standard deviation of the noise were identified using Mini Analysis software (v6.0.8, Synaptosoft, Decatur, GA). These signals were manually inspected, and artifacts or other misidentified peaks were excluded from the analysis. Double peaks, or peaks that occurred less than 1 sec after a prior peak, were manually excluded from the dataset. Peaks manually excluded accounted for less than 5% of the total number of peaks identified by the program.

### Generation of bone marrow derived mast cells (BMMCs)

Bone marrow progenitor cells were harvested from the femurs of adult males and females developmentally exposed to control, low PBDE, or high PBDE. Following established protocols^102,104^, the isolated cells were cultured in a controlled environment at 37°C with 5% CO2. The cultures were maintained in 150 cm² cell culture flasks filled with 70 mL of complete medium, which consisted of RPMI 1640 (containing L-glutamine), supplemented with 10% heat-inactivated fetal bovine serum, 1x MEM non-essential amino acids, 10 mmol/L HEPES buffer, 1 mmol/L sodium pyruvate, 100 U/mL penicillin, and 100 µg/mL streptomycin. Additionally, the medium was enriched with recombinant murine interleukin-3 (IL-3; 5 ng/mL) and stem cell factor (SCF; 5 ng/mL) from R&D Systems, Minneapolis, MN. Non-adherent cells were gently transferred to fresh complete medium every 4-5 days. 6 weeks later, purity of mast cell population (>98%) was confirmed using 0.5% toluidine blue staining at pH 0.5.

### Intracellular calcium mobilization

BMMCs were washed and seeded in calcium assay buffer at a density of 1.25×10^5^ on the day of the experiment. Intracellular calcium flux was measured using the Fluo-4 NW Calcium Assay Kit (F36206, Thermo Fisher Scientific) according to the manufacturer’s instructions. Cells were plated in black-walled 96-well plates and loaded with Fluo-4 NW dye at 37 °C for 45 minutes. Following dye loading, fluorescence was recorded at an excitation wavelength of 494 nm and emission at 516 nm at 37 °C using a Varioskan LUX multimode microplate reader (Thermo Fisher). Baseline fluorescence was recorded for 10 seconds before the addition of 60 μg/mL compound 48/80 (Sigma-Aldrich) or 20 μM ionomycin (Thermo-fisher). Fluorescence changes were monitored continuously for a total of 60 seconds to assess real-time intracellular calcium mobilization in response to stimulation.

### Beta-Hexosaminidase assay

BMMCs were seeded at a density of 1 × 10L cells per well in a clear 96-well plate in Tyrode’s buffer supplemented with 0.1% BSA (Fisher Scientific) and allowed to settle for 30 minutes at 37 °C. For IgE-mediated stimulation, cells were sensitized overnight with 0.8 µg/mL anti-DNP IgE (SPE-7, Sigma-Aldrich). On the day of the assay, cells were stimulated for 1 hour at 37 °C with 60 µg/mL compound 48/80 or vehicle control in a final volume of 100 µL Tyrode’s buffer. Following stimulation, plates were centrifuged at 1500 rpm for 5 minutes. A 30 µL aliquot of supernatant was transferred to a new 96-well plate. The remaining buffer was discarded, and cell pellets were lysed with 100 µL of 0.1% Triton X-100 in Tyrode’s buffer by pipetting up and down. A 30 µL aliquot of each lysate was transferred to the corresponding wells of the new plate. To quantify β-hexosaminidase activity, 10 µL of 3 mg/mL p-nitrophenyl-N-acetyl-β-D-glucosaminide (NAG) substrate in 0.1 M citrate buffer (pHL4.5) was added to each sample and incubated at 37L°C for 1 hour. The reaction was stopped by adding 200 µL of 0.2 M glycine buffer (pHL10.7). Absorbance was measured at 405 nm using a Varioskan LUX multimode microplate reader (Thermo Fisher Scientific). The percentage of β-hexosaminidase release was calculated as the ratio of absorbance in the supernatant to the total absorbance (supernatant + pellet), providing an index of mast cell degranulation.

### IgE-mediated stimulation of BMMCs

BMMCs were first sensitized overnight with 0.5 μg/mL of monoclonal anti-DNP IgE (clone SPE-7, Sigma-Aldrich) in complete RPMI medium. The following day, cells were washed and resuspended in fresh medium, then plated at a density of 1 × 10L cells per well in 1LmL in 12-well tissue culture plates. After a 1-hour equilibration at 37 °C in a humidified incubator with 5% CO₂, cells were stimulated with either vehicle or 15 ng/mL DNP-HSA (Sigma-Aldrich) for 30 minutes. Following stimulation, cells were harvested for cytospin analysis or transmission electron microscopy.

### Cytospin Analysis

Cytospin preparations were generated using an Epredia™ Cytospin™ 4 Cytocentrifuge by centrifuging 100 μL of cell suspension per slide at 1500 rpm for 5 minutes. Slides were allowed to air-dry and then stained with 0.5% toluidine blue in 0.5N HCl for 10 minutes to visualize mast cell granules. After rinsing and mounting, slides were imaged using a light microscope at 20× magnification. For each sample, a minimum of five non-overlapping fields were captured per slide under identical exposure settings, and images were coded to ensure blinded analysis. T-blue staining intensity, cell number, and individual cell area were quantified using ImageJ software by a blinded observer. Average optical density per cell and mean cell area were calculated from the identified mast cell population in each image.

### Transmission electron microscopy

BMMC pellets were fixed in 2.5% glutaraldehyde in 0.1LM phosphate buffer (Electron Microscopy Sciences) and stored at 4 °C. Samples were post-fixed in 1% osmium tetroxide in 0.1 M sodium phosphate buffer for 1 hour, dehydrated through a graded acetone series, and embedded in Spurr’s epoxy resin. Ultrathin sections (70 nm) were cut using a diamond knife and mounted on 200-mesh copper grids. Sections were stained with 4% aqueous uranyl acetate for 30 minutes, followed by Reynolds’ lead citrate for 14 minutes. Grids were imaged using a Talos F200X G2 Analytical Scanning Transmission Electron Microscope (Thermo Fisher Scientific) at magnifications of 500× and 3000×. Cell area, vesicle number and morphology, and mitochondrial number were quantified using ImageJ (NIH) by an observer blinded to treatment condition. Mitochondria were identified by their smaller size, double membranes, and internal cristae. Vesicles were distinguished by their larger, spherical appearance and were classified as either empty or containing electron-dense granules. A representative image showing these features is presented in **Figure 3E**.

### RNA sequencing

Unstimulated BMMCs were seeded into 12-well plates and allowed to settle for 1 hour at 37 °C and 5% CO₂. Cells were then centrifuged at 1250 rpm for 5 minutes; pellets and supernatants were separated and stored at –80 °C. Total RNA was extracted from 1.8 × 10L cells per sample using Trizol® Reagent (Invitrogen) followed by column purification with the Direct-zol RNA Microprep Kit (Zymo, R2062). RNA concentration and purity were assessed using a NanoDrop spectrophotometer; only samples with A260/280 > 1.8 and clear absorbance profiles were included. Average RNA yield was 88 ng/μL in a 30 μL final volume. RNA-seq libraries were generated and sequenced by the NCSU Genomics Core using the Illumina platform.

### RNA sequencing bioinformatic analysis

Raw reads were quality-checked using FastQC and trimmed as needed. High-quality reads were aligned to the mm39 mouse reference genome using the STAR aligner^105^. Gene-level read counts were generated with htseq-count^106^, and the resulting count matrix was analyzed in R using the DESeq2 package^107^. Genes with low counts across samples were filtered out. Differential expression was assessed within sex (PBDE vs. vehicle) using a linear model with Benjamini-Hochberg correction. Genes with adjusted p-values (padj) < 0.05 were considered significant. For each sex, differentially expressed genes were submitted to Ingenuity Pathway Analysis (IPA) to identify significantly enriched canonical pathways (–log₁₀p > 2, |z-score| > 1.6). To examine transcriptional convergence and divergence, DEGs from males and females were compared to identify overlapping genes. From this list of 669 shared DEGs, we ranked the top 50 genes regulated in the same direction (concordant) and the top 50 regulated in opposite directions (discordant) across sexes. Separate IPA analyses were performed on each list to identify pathways and upstream regulators associated with transcriptional convergence or divergence.

### Statistical analysis and figures

All statistical analyses and data visualizations—except for RNA-seq analyses—were performed using GraphPad Prism 9. An alpha level of 0.05 was used to determine statistical significance across all experiments. To compare two conditions (e.g., treatment effects within sex), we used two-tailed t-tests or Mann–Whitney tests when data did not meet assumptions of normality. For experiments assessing interactions between conditions, two or three-way ANOVAs were conducted. In studies involving repeated measures (e.g., calcium imaging, rectal temperature), we applied repeated-measures ANOVAs with treatment and time as factors. Fisher’s LSD post hoc test were used to identify significant treatments. Full statistical results and group size are provided in the corresponding figure legends. n represents either an individual mouse offspring (max 2 mice per sex per litter used to minimize liter effects), or independent cell preparations from a single mouse, as specified in each figure. Data are expressed as mean ± standard error of the mean (SEM).

## Results

### 1. Perinatal PBDE exposure leads to male-specific changes in body composition, locomotor activity, and anxiety-like and social behaviors

No differences in litter size or sex were identified between treatment groups **(Supplementary fig. 1A,B).** Body composition analyses at PN56 revealed that developmental PBDE exposure had no detectable effect on body weight or composition in females. In contrast, males exposed to either low or high doses of PBDEs exhibited a dose-dependent reduction in body weight compared to controls (**Fig. 1C**). Interestingly, this was accompanied by increased total body fat and reduced lean mass, with greater adiposity observed at the higher PBDE dose (**Figs. 1D–E**), indicating a shift in body composition toward fat accumulation despite reduced overall mass.

We next assessed whether developmental PBDE exposure impacted behavior in adulthood. Effects in the open field and social interaction tests were subtle and appeared to affect both sexes in the same direction. In the open field, although no significant differences were observed when analyzing sexes separately, a main effect of treatment emerged when sexes were combined, with high PBDE exposure associated with reduced time spent in the center of the arena (**Fig. 1F, main effect of treatment depicted with** α **symbol**), without changes in total distance traveled (**Fig. 1I**). In the social interaction test, while no differences in social vs. object investigation were detected (**Fig. 1G**), there was a main effect of treatment on time spent in the corners, with high PBDE exposure linked to increased corner time in both sexes (**Fig. 1J**). These findings suggest that developmental exposure to high-dose PBDEs induced a subtle, sex-independent increases in anxiety-like behavior in adulthood. Intriguingly, however, behavior in the elevated plus maze revealed a distinct, male-biased phenotype. Males exposed to either low or high doses of PBDEs exhibited a dose-dependent increase in time spent in the open versus closed arms (**Fig. 1H**). While this is typically interpreted as reduced anxiety-like behavior, high-dose PBDE males also showed increased average velocity in the open arms (**Fig. 1K**), which is consistent with previous studies^35^ and suggest that the effect may instead reflect increased locomotor drive or behavioral disinhibition, rather than true anxiolysis.

### 2. Perinatal PBDE exposure impairs mast cell responsiveness to Fc**ε**RI-and MrgprB2-mediated activation in adulthood in both males and females

To evaluate the effects of developmental PBDE exposure on mast cell function, we employed a well-established *in vivo* model of IgE-mediated passive systemic anaphylaxis (PSA) to assess FcεRI-dependent mast cell activation—one of the most widely studied pathways in the context of mast cell–mediated allergic responses^108,109^. In this model, systemic mast cell degranulation-and particularly the release of histamine and chymase^71^-triggers a rapid drop in core body temperature, making hypothermia a reliable and quantifiable readout of mast cell activation^102,103,110^. Adult male and female offspring were sensitized via intraperitoneal (i.p.) injection with 5 μg of anti-DNP IgE monoclonal antibody, followed 24 hours later by an i.p. challenge with 50 μg of DNP to induce anaphylaxis, as previously described^111^. Rectal temperature was recorded every 5 minutes following DNP administration to measure PSA-induced hypothermia. At 30 minutes post-challenge, mice were euthanized, and blood, mesentery, and meninges were collected for downstream analysis of plasma histamine levels and mast cell degranulation by histology (**Fig. 2A**). The mesentery and meninges were selected for mast cell histological analysis due to their high mast cell density, tissue accessibility, and relevance to both immune and neuroimmune responses. Mesenteric mast cells are abundant, tissue-resident populations positioned along the gastrointestinal tract—a primary site of antigen exposure—and are commonly examined in the context of PSA^102^. The meninges, specifically the dura mater, are another site of abundant mast cell accumulation. Meningeal mast cells are increasingly recognized for their role in neuroimmune regulation, and our previous work demonstrated that this population is sensitive to developmental stress exposure^112^.

**Figure 2.**
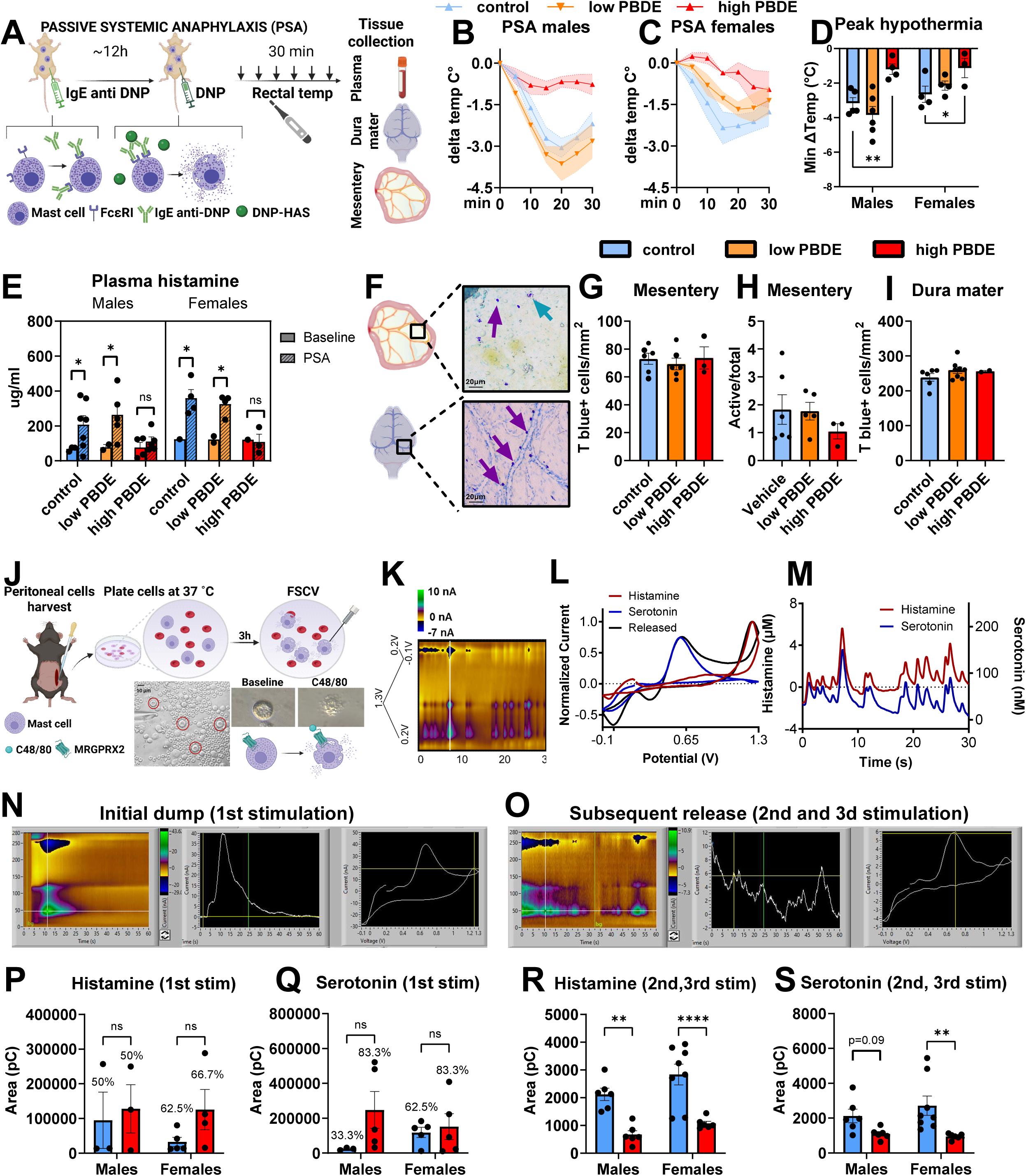
Developmental PBDE exposure blunts stimulus-induced release of histamine in adult tissue-resident mast cells. **(A)** Experimental timeline for passive systemic anaphylaxis (PSA) testing **(B)** Rectal temperature change over time in males (n = 4-6/group). Two-way repeated-measures ANOVA revealed significant main effects of time (F (1.772, 21.27) = 36.21, *p* < 0.0001), treatment (F (2, 12) = 8.562, *p* = 0.0049), and a time × treatment interaction (F (12, 72) = 4.693, *p* < 0.0001). Fisher’s LSD showed significantly blunted hypothermia in high PBDE vs. control at 15 min (*p* = 0.0069), 20 min (*p* = 0.0009), and 25 min (*p* = 0.0009), and a more modest difference at 30 min (*p* = 0.0414) **(C)** Rectal temperature change over time in females (n = 4/group). Two-way repeated-measures ANOVA detected main effects of time (F (1.601, 12.81) = 10.14, *p* = 0.0034) and treatment (F (2, 8) = 6.812, *p* = 0.0187), but no significant interaction. Fisher’s LSD comparisons showed reduced temperature drop in high PBDE vs. control at 5 min (*p* = 0.0463), 15 min (*p* = 0.0411), and 20 min (*p* = 0.0160). **(D)** Peak drop in rectal temperature following PSA challenge in males (n = 4-6/group). Two-way ANOVA showed a main effect of sex (F (1, 20) = 4.6, *p* = 0.004), and PBDE treatment (F (2, 20) = 10.38, *p* = 0.0008). Fisher’s LSD post hoc test showed that in both, males (p=0.0034) and females (p=0.03), high PBDE blunted hypothermic responses compared to control treatment. **(E)** Plasma histamine levels (n = 2-7/group). Three-way ANOVA showed a main effect of PSA (F (1, 20) = 8.9, *p* = 0.0007). Fisher’s LSD revealed that, while, compared to baseline, PSA induced an increase in plasma histamine in control males (p=0.04) and females (p=0.03), as well as low PBDE males (p=0.03) and females (p=0.04), this was not true in high PBDE males or females. **(F)** Representative images of toluidine blue–stained mast cells in the mesentery and dura mater. Purple arrows indicate inactive mast cells (dense T blue staining), turquoise arrows indicate actively degranulating mast cells. **(G)** Quantification of mesenteric mast cell density (n = 3-7/group). No significant differences were observed across groups. **(H)** Quantification of mesenteric mast cell activation levels in mesentery (n = 3-7/group). No significant differences were observed across groups. **(I)** Quantification of dural mast cell density (n = 2-6/group). No significant differences were observed across groups. **(J)** Schematic of peritoneal mast cell isolation and recording of histamine release using fast-scan cyclic voltammetry following compound 48/80 stimulation. **(K)** Representative color plot containing 30 sec of raw voltammetric data collected upon degranulation of a single mast cell. Voltage is plotted on the Y axis, time is plotted on the x axis, and current is represented in color. **L)** Normalized CVs for serotonin and histamine standards directly overlaid with a CV recorded ∼7 sec after stimulation (white line in (K)). **M)** Concentration vs time traces reflecting serotonin and histamine dynamics, corresponding to the data shown in (K). The correlated signals suggest co-release. **(N)** Representative color plot of raw voltammetric data collected upon first stimulation with C48/80. **(O)** Representative color plot of raw voltammetric data collected upon 2^nd^ and 3rd stimulations with C48/80. **(P)** Total histamine release per mast cell in males and females (n = 6/group, only a subset showed initial dump, depicted in % above individual bars) upon first stimulation with C48/80. No statistically significant differences were seen between groups. **(Q)** Total serotonin release per mast cell in males and females (n = 6/group, only a subset showed initial dump, depicted in % above individual bars) upon first stimulation with C48/80. No statistically significant differences were seen between groups, although a higher percent of cells in high PBDE males showed initial dump compared to controls **(R)** Histamine release per mast cell in males and females upon subsequent stimulations with C48/80. Two-way ANOVA showed a main effect of sex (F (1, 22) = 4.3, *p* = 0.0499) and stimulation (F (1, 22) = 35.91, *p* < 0.0001). Fisher’s LSD revealed that, compared to controls, high PBDE males (p=0.001) and females (p<0.0001) showed significantly reduced histamine release. **(S)** Serotonin release per mast cell in males and females upon subsequent stimulations with C48/80. Two-way ANOVA showed a main effect of stimulation (F (1, 22) = 12.42, *p* = 0.002). Fisher’s LSD revealed that, compared to controls, high PBDE females (p=0.004) showed significantly reduced serotonin release, while males showed only a trend (p=0.09).

**Figure 3.**
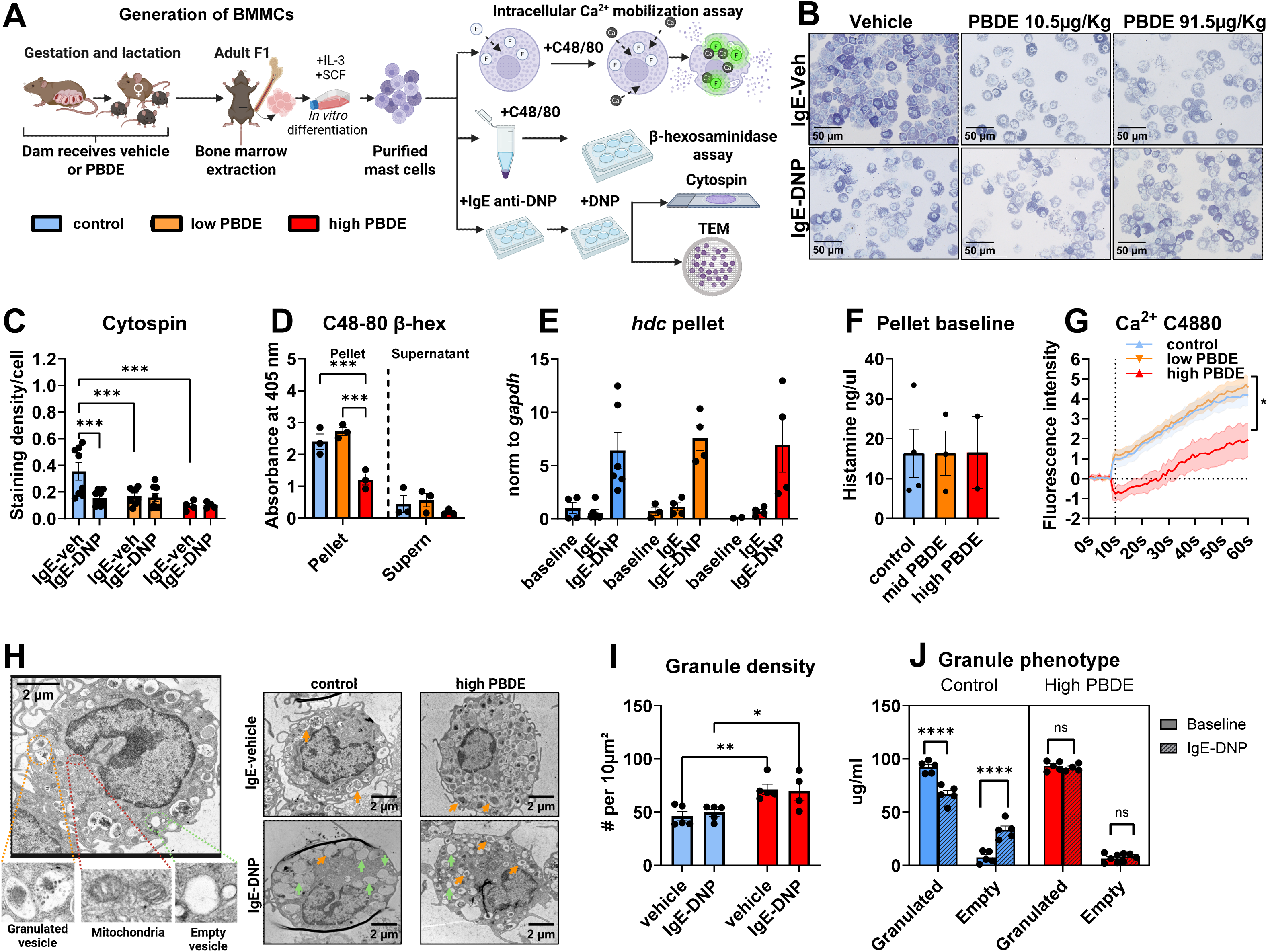
BMMCs derived from adult animals developmentally exposed to PBDE show impaired granule phenotype and stimulus-induced calcium mobilization, without showing differences in overall histamine synthesis or storage. **(A)** Generation of BMMCs and experimental design. **(B)** Toluidine blue staining of IgE-sensitized BMMCs after vehicle or DNP exposure. **(C)** T blue staining density quantification (n = 4-8/group). Two-way ANOVA showed a main effect of PBDE exposure (F (2, 34) = 7.106, *p* = 0.002), DNP stimulation (F (1, 34) = 4.345, p=0.04) and interaction (F (2, 34) = 4.527, p=0.02). Fisher LSD showed that only in control BMMCs DNP stimulation reduced staining density compared to vehicle (p=0.0003). Additionally, both low (p=0.0008) and high PBDE (p=0.0002) BMMCs showed reduced staining density at baseline compared to controls. **(D)** β-hexosaminidase assay (n = 3/group). Two-way ANOVA showed a main effect of PBDE exposure (F (2, 12) = 13.55, *p* = 0.0008), C48/80 stimulation (F (1, 12) = 122.3, p<0.0001) and interaction (F (2, 12) = 5.129, p=0.02). Fisher LSD showed that high PBDE compared to both controls (p=0.0008) and low PBDE (p=0.0001) showed reduced β-hexosaminidase pellet content. (E) *Hdc* expression measured by qPCR in BMMC pellets (n=4-6/group). Two-way ANOVA revealed a main effect of IgE-DNP stimulation (F (2, 28) = 24.20, *p* < 0.0001). Planned comparisons revealed that IgE-DNP elicited a comparable induction of *Hdc* across treatment groups (baseline vs IgE-DNP all p<0.002). **(F)** Histamine levels measured in BMMC pellets at baseline (no stimulation) (n=2-4/group). No differences were observed. **(G)** Calcium mobilization following C48/80 stimulation. Two-way repeated-measures ANOVA revealed significant main effects of time (F (1.957, 115.4) = 55.67, *p* < 0.0001), PBDE treatment (F (2, 59) = 3.788, *p* = 0.002), and a time × treatment interaction (F (120, 3540) = 2.167, *p* < 0.0001). Fisher’s LSD post hoc comparisons showed significantly blunted Calcium mobilization in high PBDE vs. control at every timepoint starting immediately after C48/80 addition (p=0.002-0.003) **(H)** Representative transmission electron microscopy images of IgE-sensitized BMMCs after vehicle or DNP challenge. Orange arrows show granules with electron-dense content, green arrows show empty granules **(I)** Granule density (number of granules per cell, n=5/group). Two-way ANOVA revealed a main effect of PBDE exposure (F (1, 16) = 20.11, p=0.0004), and DNP stimulation (F (1, 16) = 4.687, p=0.04). Fisher LSD revealed that high PBDE BMMCs showed increased granule density compared to controls (p=0.0005). **(J)** Granule phenotype (empty vs. granulated vesicle ratio, n=5/group). Three-way ANOVA showed an interaction effect between vesicle type, PBDE treatment, and stimulation (F (1, 30) = 37.61, p<0.0001). Fisher LSD revealed that only control BMMCs showed an increase of empty vesicles after DNP challenge (p <0.0001). **(K)** Mitochondrial density (mitochondria per cell area) in BMMCs after IgE sensitization and DNP challenge.n=5/group). Only a significant interaction effect was detected in two way ANOVA (F (1, 16) = 5.690, p=0.03). Fisher LSD revealed that control BMMCs showed a reduction in dense (p<0.0001), as well as increase in empty granules (p<0.0001) after DNP stimulation, which was not observed in high PBDE BMMCs.

Our results showed that, while control and low-PBDE–exposed animals exhibited the expected hypothermic response to PSA, this response was markedly blunted in males and females developmentally exposed to high PBDE (**Figs. 2B–D**). This outcome was supported by plasma analyses: compared to non-challenged animals, IgE-DNP significantly increased plasma histamine levels at 30 min in control and low-PBDE groups, but no such increase was observed in high-PBDE–exposed animals of either sex (**Figs. 2E**). Notably, tissue-resident mast cell density or activation levels, as measured by T blue staining of the intestinal mesentery windows and dura mater, did not differ across exposure groups (**Figs. 2F–I**, male and female data pooled), indicating that the blunted physiological and biochemical responses were not due to reduced mast cell numbers or activation levels.

Next, to further characterize mast cell phenotypes in PBDE-exposed animals and determine whether the functional deficits extended beyond FcεRI signaling, we isolated peritoneal mast cells and used fast-scan cyclic voltammetry to assess histamine and serotonin release in response to C48/80 (**Fig. 2J–M**). C48/80 triggers mast cell degranulation by activating MrgprB2 (the mouse homolog of human MRGPRX2), a G protein–coupled receptor involved in host defense and immune modulation, as well as in the pathogenesis of pseudo-allergic drug reactions, pruritus, pain, and inflammatory diseases^113,114^. Our results showed that, in control animals, roughly half to two-thirds of mast cells exhibited a large, monophasic “initial dump” of amine release in response to the first stimulation (**Fig. 2N**). While this response was modestly more frequent in PBDE-exposed cells (percentages shown above bars in **Fig. 2P,Q**), the total charge released during this initial burst did not differ significantly between groups. In contrast, histamine and serotonin release during the second and third stimulations were significantly reduced in PBDE-exposed cells of both sexes (**Figs. 2O,R,S**), indicating that developmental PBDE exposure impairs the ability of mast cells to sustain amine release across repeated stimuli, which could have important physiological implications. Because histamine and serotonin are both cleared from extracellular fluid within minutes^115,116^, a single burst that is not followed by continued release will result in lower circulating levels over time. In the PSA model, mast cells are continually stimulated over an extended period by circulating antigen that crosslinks IgE on their surface, promoting sustained degranulation *in vivo*. Thus, these results suggest that the impaired ability to maintain amine release in PBDE-exposed cells contributes to the reduced plasma histamine levels and attenuated hypothermic response observed 30 min following PSA. Intriguingly, while the total cumulative release of histamine (initial dump plus subsequent events) did not differ between groups (**Supplementary fig. 1C**), the cumulative release of serotonin was increased in PBDE exposed males but not females (**Supplementary fig. 1D**). Mast cell derived serotonin has been recently implicated in promoting fat storage^66^, potentially explaining the increased fat % observed in PBDE exposed males.

### 3. Mast cells derived from adult animals developmentally exposed to PBDE show impaired granule phenotype and stimulus-induced calcium mobilization, without showing differences in overall synthesis storage

To further determine whether the effects of developmental PBDE exposure on mast cell function were driven by the tissue microenvironment—which plays a major role in shaping mature mast cell phenotype^117–120^—or by intrinsic reprogramming of mast cell progenitors, we isolated bone marrow—the primary source of mast cell precursors during the postnatal period^121,122^—from adult offspring and differentiated bone marrow– derived mast cells (BMMCs) *in vitro* using established protocols^102,123^ (**Fig. 3A**). Consistent with *in vivo* findings, no sex-dependent effects were observed across any of the BMMC outcomes assessed; therefore, data from males and females were pooled for all subsequent analyses, with the exception of electron microscopy (EM), which was performed only in male-derived cells.

To characterize BMMC phenotypes, we first used toluidine blue staining of IgE-sensitized BMMCs (**Fig. 3B**). Toluidine blue is a metachromatic dye that binds to acidic proteoglycans such as heparin and chondroitin sulfate^124,125^, which are enriched in mature mast cell granules and thus commonly used to identify mast cells. BMMCs derived from both control and PBDE-exposed animals showed positive metachromatic staining, confirming mast cell identity. However, IgE-sensitized BMMCs from low and high PBDE-exposed animals exhibited reduced staining intensity compared to controls (**Fig. 3C**). Additionally, while control BMMCs showed a clear reduction in staining density following DNP stimulation—consistent with granule release—this reduction was not observed in PBDE BMMCs. These findings suggest that mast cells derived from PBDE exposed animals may have reduced granule density or altered granule composition following IgE sensitization and fail to undergo typical granule discharge upon activation.

To test this interpretation and determine whether these deficits extended to other canonical measures of mast cell function, we assessed β-hexosaminidase (β-hex) release following stimulation with C48/80. β-hex is a lysosomal enzyme stored in mast cell granules and released upon activation; measuring its levels in the supernatant and pellet provides insight into both granule content and degranulation efficiency^126^. Contrary to expectations, the proportion of β-hex released (supernatant/total) following C48/80 stimulation did not differ significantly between groups. However, a closer examination revealed that BMMCs from high PBDE–exposed animals exhibited significantly lower total β-hex content at baseline compared to both controls and the low PBDE group (**Fig. 3D**). This was also true for Ionomycin-induced mast cell degranulation (**Supplementary fig. 1E**). These results suggest that although release ratio is preserved, the overall amount of β-hex released is reduced due to diminished granule content or maturity, aligning with the toluidine blue results.

Building on these findings, and our observations of blunted sustained histamine release following IgE-DNP and C48/80 stimulation in tissue-resident mast cells, we next hypothesized that developmental PBDE exposure might impair histamine synthesis and/or overall storage. However, neither mRNA expression of histidine decarboxylase (Hdc)—the rate-limiting enzyme for histamine synthesis—at baseline or following IgE-DNP stimulation (**Fig. 3E**), nor total baseline histamine content (**Fig. 3F**), differed between exposure groups, suggesting that impaired histamine release is not due to reduced histamine synthesis or whole-cell storage capacity.

Lastly, because calcium influx is required for granule trafficking, membrane fusion, and sustained mediator release in mast cells^127,128^, we measured C48/80-induced calcium mobilization in BMMCs. Intriguingly, calcium mobilization was significantly blunted in BMMCs from the high PBDE group compared to controls (**Fig. 3G**). Because our calcium assay records the average fluorescence across a population of cells, it reliably captures bulk cytosolic calcium elevations but cannot detect spatially restricted calcium “puffs”— localized, transient increases in calcium that occur near the plasma membrane^127,129^. These puffs are often the first step in mast cell activation, initiating granule fusion by triggering exocytosis of the readily releasable pool. In individual mast cells, these localized signals typically propagate into broader calcium waves and oscillations that spread throughout the cytoplasm. This propagation is necessary to sustain granule trafficking and fusion over time, supporting extended mediator release. Under this framework, mast cells from high PBDE animals may still generate initial calcium puffs sufficient for a first burst of histamine release, but fail to achieve the more global and sustained calcium dynamics needed to maintain secretion beyond the initial response. Together, these results indicate that the deficits in sustained histamine release observed in PBDE-exposed tissue-resident mast cells may reflect disruptions in granule packaging, positioning, or in the calcium-dependent signaling required to support extended degranulation.

To explore this possibility at the ultrastructural level, we performed electron microscopy on IgE-sensitized BMMCs at baseline and 1h following IgE-DNP stimulation. Analysis of electron microscopy images revealed that, although BMMCs from both control and PBDE-exposed animals displayed typical mast cell morphology—including a large central nucleus and numerous cytoplasmic granules (**Fig. 3H**)—BMMCs from PBDE-exposed animals exhibited a significantly higher granule density, measured as the number of granules per cell (**Fig. 3I**). This increase in granule number may reflect impaired granule maturation, as immature granules accumulate when fusion and processing steps are disrupted^130,131^. Following IgE-DNP stimulation, control BMMCs showed a clear increase in the proportion of empty granules—consistent with active degranulation—whereas this response was absent in high PBDE–exposed BMMCs (**Fig. 3J**). Taken together, these findings suggest that PBDE-exposed mast cells exhibit both defective granule maturation and a failure to mobilize and release granules during activation, potentially explaining their impaired sustained secretory function.

### 4. Developmental PBDE exposure induces sex-specific transcriptional phenotypes in BMMCs

To investigate the molecular mechanisms underlying the blunted mast cell responses observed after developmental PBDE exposure, we performed RNA sequencing on BMMCs from adult males and females exposed perinatally to either control vehicle or high-dose PBDE (**Fig. 4A**). Despite no measurable sex differences in mast cell function across *in vivo*, *ex vivo*, and *in vitro* assays, transcriptomic analysis revealed marked sex-specific responses. Of the differentially expressed genes (DEGs) identified between PBDE and vehicle conditions, only ∼17% (669 of 3834) overlapped between males and females (**Fig. 4B**), indicating limited transcriptional convergence.

**Figure 4.**
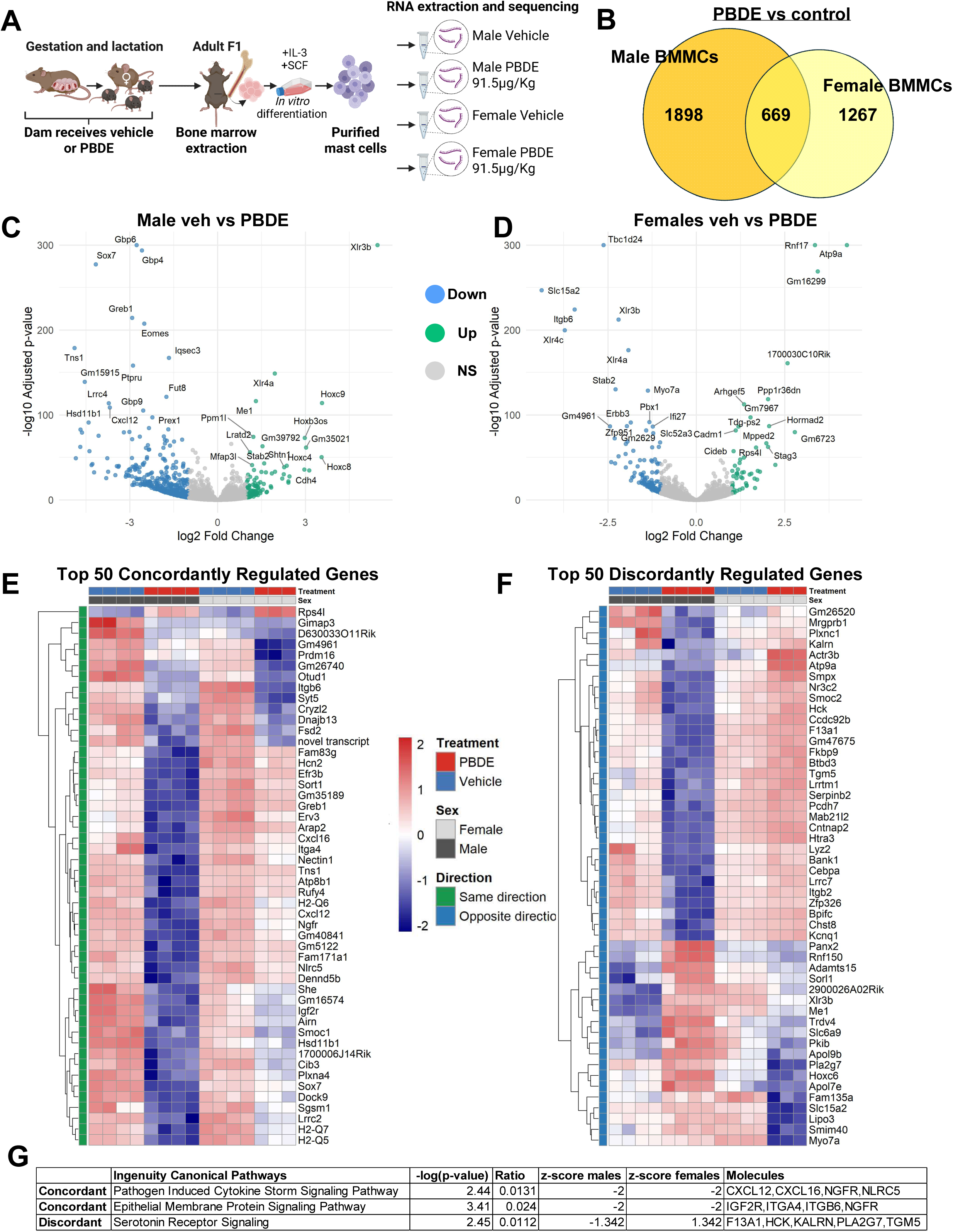
Developmental PBDE exposure induces sex-specific transcriptional signatures in mast cells. **(A)** Experimental design RNA sequencing of BMMCs (n = 4/sex/treatment). **(B)** Venn diagram showing differentially expressed genes (DEGs) in PBDE-exposed vs. vehicle BMMCs in males and females. **(C–D)** Volcano plots of DEGs in males **(C)** and females **(D)**, comparing PBDE vs. vehicle. Colored points indicate significantly upregulated (green) and downregulated (blue) transcripts (adjusted p < 0.05, log2 fold change > 1 or < –1); grey points represent non-significant genes. **(E)** Heatmap of the top 50 concordantly regulated genes across sexes, defined as genes significantly upregulated or downregulated by PBDE exposure in both males and females. Rows represent genes; columns represent individual animals. Color scale reflects log2-transformed, z-score–normalized expression. **(F)** Heatmap of the top 50 discordantly regulated genes, defined as genes significantly upregulated in one sex and downregulated in the other. Sex, treatment, and directionality are color-coded in the top and left annotations. **(G)** Ingenuity Pathway Analysis (IPA) of canonical pathways enriched among concordantly (top 2) and discordantly (top 1) regulated genes. Columns indicate pathway name, –log10(p-value), gene ratio, activation z-score in males and females, and representative molecules. Negative z-scores indicate pathway inhibition; positive scores indicate activation.

Volcano plots illustrate distinct DEG profiles in males (**Fig. 4C**) and females (**Fig. 4D**). To assess the functional implications of these sex-specific transcriptomic changes, we performed Ingenuity Pathway Analysis (IPA) separately for each sex (**Supplementary figs 2A,B**; displaying pathways with –log₁₀pL>L2 and |z-score|L>L1.6; full results in **Supplementary Tables 1 and 2**). Despite minimal overlap in DEGs between sexes, IPA identified three canonical pathways that were significantly altered in both males and females: *Tuberculosis Active Signaling* was upregulated, while *FAK Signaling* and *Pathogen-Induced Cytokine Storm Signaling* were downregulated in both sexes. However, closer examination revealed that the specific genes driving these pathway enrichments were largely sex-specific (**supplementary fig 2C**). For instance, no individual genes overlapped between sexes in the *Tuberculosis Active Signaling pathway*. In contrast, *FAK Signaling* and *Pathogen-Induced Cytokine Storm Signaling* showed partial overlap. Notably, shared genes included CCR1, which mediates mast cell activation, migration, and degranulation^132–134^ and FCER1G, a critical subunit of the high-affinity IgE receptor^135,136^. These shared components suggest that, despite broader transcriptomic divergence, PBDE-induced mast cell impaired mediator release in males and females may arise from convergent disruption of key effector mechanisms.

To better understand the molecular basis of the shared mast cell dysfunction observed in both sexes, we next examined whether transcriptional convergence could explain this functional similarity. Specifically, we identified genes that were regulated in the same or opposite directions in males and females. We ranked the top 50 concordantly regulated genes—those similarly up-or downregulated in both sexes (**Fig. 4E; Supplementary Table 3**)—and the top 50 discordantly regulated genes, which changed in opposite directions (**Fig. 4F; Supplementary Table 4**). We then performed separate IPA for each gene set, using input files that included gene symbols, log₂ fold changes, and adjusted p-values for each sex. This strategy enabled a direct comparison of pathways and upstream regulators associated with transcriptional convergence versus divergence.

Despite broader transcriptomic differences, IPA revealed that concordantly regulated genes were enriched for immune-related pathways that were downregulated in both sexes (**Fig. 4G**). Notably, the *Epithelial Membrane Protein Signaling pathway* was downregulated (–log₁₀p = 3.41, z-score = –2), implicating disruption of granule biogenesis, composition, and trafficking through genes like IGF2R—whose major function is the trafficking of lysosomal enzymes to endosomes and their subsequent transfer to lysosomes^137^—ITGA4, a cell surface adhesion molecule involved in mast cell maturation¹³L, ITGB6, which regulates mast cell protease expression^138^, and NGFR, the receptor for nerve growth factor that contributes to mast cell development and differentiation^139,140^. These alterations could potentially explain the increased granule density and reduced staining intensity observed in PBDE-derived mast cells. Interestingly, downregulated IGF2R, part of the insulin receptor family, could also contribute to the blunted calcium mobilization observed in PBDE BMMCs. Receptors in the insulin receptor family are known to strongly activate the PI3K pathway^141^. This pathway enhances extracellular calcium influx—a critical second phase of calcium signaling following activation-induced calcium release from intracellular stores—and also contributes to Protein Kinase C activation^136^, both of which are required to sustain granule trafficking and exocytosis.

Additionally, the *Pathogen-Induced Cytokine Storm Signaling pathway* (–log₁₀p = 2.44, z-score = –2 in both sexes) was suppressed through partially overlapping genes such as CXCL12, CXCL16, NGFR, and NLRC5, suggesting a shared dampening of cytokine and chemokine signaling. Given that cytokine signaling supports mast cell priming and communication with neighboring immune cells^64,142,143^, this broad suppression may reflect a state of immune hypo-responsiveness in PBDE-exposed mast cells.

In contrast, analysis of discordantly regulated genes uncovered that the *Serotonin Receptor Signaling* pathway was significantly altered (–log₁₀p = 2.45), but in opposite directions in males (z = –1.34) and females (z = +1.34). While this pathway is classically associated with neuromodulation, several of the contributing genes—F13A1, HCK, KALRN, and PLA2G7—are functionally linked to mast cell activation, granule maturation, and/or cytoskeletal reorganization^144–147^, raising the possibility that mast cell responses in contexts beyond degranulation via IgE-FcεRI or MRGPRX2 IgE could be sex-specifically modulated by PBDEs, a hypothesis that warrants further investigation.

## Discussion

Human and animal exposure to PBDEs remains widespread despite global restrictions on their production and use. This is particularly concerning for developing individuals, who are exposed through placental transfer and lactation during periods of heightened vulnerability due to immature detoxification systems and ongoing development of the immune, nervous, and endocrine systems^148^. While developmental PBDE exposure has been linked to a wide range of multisystemic outcomes—from endocrine^34,36–38^ and reproductive impairments^35,39^ to metabolism^40–43^ and neurobehavioral deficits^44–46^—its effects on immune function remain comparatively underexplored. This represents a critical gap, particularly given growing evidence that the immune system not only shapes the early development of other physiological systems but also maintains continuous, bidirectional communication with them throughout life^149–153^. As such, immune dysfunction could contribute to, or even drive, some of the systemic effects attributed to early-life PBDE exposure.

To explore immune contributions to PBDE-induced dysfunction, we focused on mast cells. Mast cells are among the earliest immune cells to mature and establish residence in tissues during development^154^. These long-lived cells are highly sensitive to environmental cues—including xenobiotics, hormones, stress, and pathogens^53–55,60–63,155,156^—and exert broad regulatory influence across immune, vascular, and neuroendocrine systems^64–71^. These properties make them a strong candidate for mediating the long-term, multisystem effects of early-life toxicant exposure.

Using human-relevant doses of PBDEs administered to dams throughout pregnancy and lactation, we found that daily developmental exposure via maternal transfer to ∼87 μg/kg—aligned with the lower end of doses used in previous mouse studies assessing metabolic and neurobehavioral outcomes^95–97^ and within 10-fold of PBDE levels measured in human serum and placenta^93,94^—leads to persistent dysfunction in mast cell release of amines in adult male and female offspring in response to both FcεRI-and MRGPRX2-mediated stimulation. These functional deficits could not be explained by differences in tissue-resident mast cell numbers. Studies using bone marrow–derived mast cells (BMMCs) from adult offspring revealed that PBDE exposure did not impair histamine synthesis or storage. Instead, we observed deficits in granule maturation and in stimulus-induced calcium mobilization—processes required for extended degranulation. RNA sequencing data of BMMCs revealed that these impairments may be driven by the downregulation of genes such as IGF2R, ITGA4, ITGB6, and NGFR, which are involved in granule composition, trafficking, and mast cell development. Importantly, because the bone marrow is the primary postnatal source of mast cells^157,158^, these findings suggest that the effects of developmental PBDE exposure on adult mast cell function are due to early reprogramming at the progenitor level, with potential implications across tissue mast cells.

In sum, to our knowledge, this is the first study to assess the effects of developmental PBDE exposure on adult mast cell physiology. Our results suggest that mast cell hypofunction may represent a previously unrecognized mechanism contributing to the long-term physiological and behavioral consequences of early-life toxicant exposure.

### Persistent impairment of sustained mast cell–derived histamine release as a potential mechanism by which developmental PBDE exposure contributes to long-term physiological and behavioral alterations

A key outcome of this study is the persistent impairment of stimulus-induced mast cell release of prestored histamine. Histamine is a biogenic monoamine with pleiotropic effects across physiological systems^72^, mediated through four distinct G protein–coupled receptors (H1R–H4R) that are differentially expressed across tissues. While several cell types can synthesize histamine, mast cells are the only tissue-resident immune cells that store it in large quantities, enabling both rapid release in response to stimuli as well as sustained, piecemeal degranulation that may exert continuous local influence^67,159^. Given histamine’s widespread actions, defects in mast cell histamine release could plausibly contribute to many of the long-term physiological and behavioral effects associated with developmental PBDE exposure.

For instance, PBDEs have been linked to altered reproductive endpoints in both sexes, including disrupted gonadal development, gametogenesis and hormone secretion^35,38,160–162^, as well as adverse pregnancy outcomes^163,164^. Mast cells are enriched in gonadal and nongonadal reproductive tissues, and increasing evidence indicates that histamine—acting through H1R, H2R, and H4R—can promote Leydig cell proliferation, steroidogenesis, and sperm viability^165–167^. It is also believed to play a role in implantation and early placental development by facilitating local tissue remodeling^168,169^.

Similarly, the effects of developmental PBDE exposure on neurodevelopment and behavior—including deficits in learning, hyperactivity, social behaviors, and increased anxiety^44,170–173^—could be at least partly mediated by altered histaminergic signaling. In the central nervous system, histamine plays key roles in modulating arousal, cognition, stress responses, and social behaviors^174–177^. Mast cells, which accumulate in meninges and brain regions such as the hypothalamus, thalamus, and hippocampus ^178–180^, not only contribute approximately 50% of total brain histamine^86,87^, but may also regulate activity of histaminergic neurons via the inhibitory H3 autoreceptor^181^—though this remains to be directly demonstrated. While the specific roles of mast cell–derived histamine in the brain are still being elucidated, accumulating evidence suggests it contributes to multiple neurobiological processes, including arousal^88^, modulation of anxiety^87,88^ and stress responses^91,92^, and organization of sex-specific neural circuits^90^. As such, persistent disruption in mast cell histamine release could contribute to the neurobehavioral phenotypes observed following developmental PBDE exposure.

Interestingly, however, excessive mast cell histamine release can also have detrimental effects. For example, histamine is a central mediator of pathological allergies^109,182,183^, contributes to increased gut inflammation and permeability^184,185^, migraines^186,187^, and can exacerbate neuroinflammation^174,188^. Therefore, it would be important to assess whether the effects of developmental PBDE exposure might also reduce susceptibility to disorders associated with excessive histamine signaling.

### Sex-specific effects and mast cell functions beyond histamine release

While this study highlights persistent defects in mast cell–derived histamine release driven by early reprogramming, our transcriptomic data reveal a broader and highly sex-specific pattern of disruption, suggesting that developmental PBDE exposure may impair additional mast cell functions beyond histamine release, contributing to sex-specific vulnerabilities. For example, in female BMMCs, the Th2 signaling pathway was selectively downregulated, marked by reduced expression of key regulators such as GATA3 and IL4. Th2 immunity—driven by interactions among mast cells, Th2 T cells, IgE-producing B cells, and dendritic cells— plays essential roles not only in defense against helminths and venoms^189–191^, but also in promoting tissue repair^192^ and contributing to protective avoidance behaviors^193,194^. Mast cells support these responses, at least in part, by producing IL-4 and histamine, which influence dendritic cell polarization, Th2 differentiation, and IgE class switching^195^. Thus, suppression of this pathway may increase susceptibility to parasitic infection, impair tissue repair, or weaken epithelial defenses in PBDE-exposed females.

Similarly, males developmentally exposed to PBDEs exhibited downregulation of the antigen presentation pathway, including reduced expression of B2M, TAPBP, and HLA-A—key components of the MHC class I pathway ^196–198^ responsible for presenting endogenous peptides to CD8⁺ cytotoxic T cells—as well as CIITA, HLA-DMA, and HLA-DOB, which are involved in the MHC class II pathway^199,200^ and facilitate the presentation of extracellular antigens to CD4⁺ T cells. Although mast cells are not classically considered antigen-presenting cells (APCs), an increasing body of evidence indicates that they can function as atypical APCs, capable of presenting antigens to both CD4⁺ and CD8⁺ T cells^201,202^. Antigen presentation is a key mechanism for shaping adaptive immune responses, and while multiple cell types can serve this role— introducing some functional redundancy^203^—mast cells may provide nonredundant, context-specific APC functions, particularly at tissue interfaces or during inflammation^204^. As such, it will be important to determine whether adaptive immune responses are altered in PBDE-exposed males, and whether these changes are driven by mast cell-specific deficiencies or broader immune dysregulation.

### Broader implications for immune programming by PBDEs

While this study focused specifically on mast cells, the fact that these alterations were evident in BMMCs—differentiated *in vitro* from hematopoietic progenitors—suggests that other hematopoietic cell lineages may also be affected by early-life PBDE exposure. Supporting this hypothesis, previous studies have reported systemic immunosuppressive effects following developmental PBDE exposure. In humans, elevated PBDE levels have been associated with reduced immune function, but only in children with autism spectrum disorder^205^, suggesting that underlying genetic or developmental susceptibilities may modulate risk. In animal models, PBDE exposure has been linked to decreased numbers of white blood cells, neutrophils, and lymphocytes in offspring^206^. Similar findings have emerged from *in vitro* studies using immune cells from marine mammals, such as harbor seals, which exhibited PBDE-induced immunosuppression^207^. Collectively, these data raise the possibility that PBDEs broadly impair immune development through effects on hematopoietic programming, potentially disrupting the functional maturation of multiple immune lineages. Future studies should investigate the epigenetic and transcriptional mechanisms driving these persistent immune alterations and determine whether mast cells function as an early sentinel for broader hematopoietic vulnerability to environmental toxicants.

### Summary and future directions

In sum, this study provides the first evidence that developmental exposure to PBDEs induces long-lasting impairments in mast cell function, characterized by reduced stimulus-induced histamine release and broader, sex-specific transcriptional reprogramming. These findings suggest a previously unrecognized mechanism by which early-life exposure to environmental toxicants could contribute to persistent physiological and behavioral dysfunctions—through immune-specific pathways. However, several limitations remain. First, the specific PBDE congener(s) responsible for these effects have not been isolated, making it difficult to determine structure–function relationships. Second, although we observed clear deficits in histamine release, other critical mast cell functions—such as synthesis of lipid mediators, cytokines, or proteases—were not directly assessed. Third, our findings do not establish whether mast cells are causal mediators of long-term PBDE-induced pathophysiology. Future studies should aim to test the sufficiency and necessity of mast cells using in vivo models that allow for mast cell-specific ablation or functional restoration. Moreover, targeted investigations into the epigenetic alterations that underlie persistent mast cell hypofunction could reveal therapeutic windows to reverse or mitigate the effects of early-life PBDE exposure.

## Acknowledgments

NDW and DDJ were supported by the Center of Human Health and the Environment grant P30ES025128. HMS and HP were supported by a grant from the National Institute of Environmental Health Sciences (R01 ES031419).

## Author contributions

Natalia Duque-Wilckens conceived the study, designed all experiments except voltammetry, contributed to in vivo procedures, performed analysis of the RNA-seq data (following initial raw data processing by Dereje Jima), and wrote the manuscript. Jared Franges conducted all in vivo mast cell stimulation experiments and in vitro BMMC assays. Lauren Malinowski performed the behavioral testing. Chathuri De Alwis and Gregory McCarthy conducted and analyzed voltammetry experiments under the guidance of Leslie Sombers. Taylor Doolittle collected body composition data. Hannahleeh Dixon and Yang Tang assisted with BMMC experiments. Helen Watson analyzed toluidine blue–stained BMMC images, and Jasmine Peace analyzed electron microscopy images. Dereje Jima performed the initial RNA-seq data processing. Heather Patisaul provided early input on PBDE exposure design, and Heather Stapleton’s lab prepared the PBDE solutions. All coauthors contributed with manuscript editing.

---

**Supplementary figure 1.**
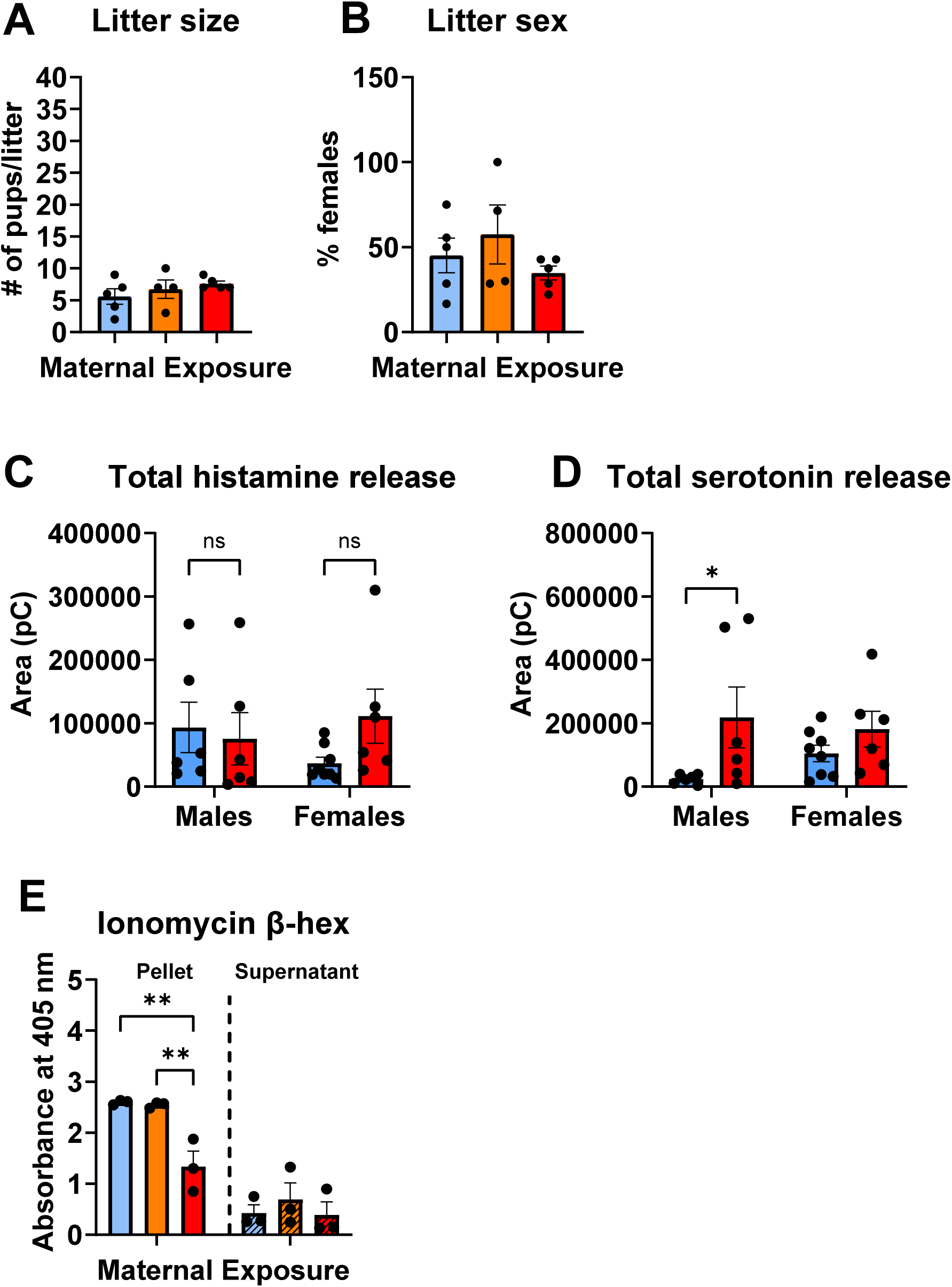
(A) Number of pups per litter. No differences were observed between treatment groups. **(B)** Percent of females in each litter. No differences were observed between treatment groups. **(C)** Cumulative histamine release per mast cell in males and females (n = 6/group) across stimulations with C48/80 as measured by voltammetry. No differences were seen between treatment groups. **(D)** Cumulative serotonin release per mast cell in males and females (n = 6/group) across stimulations with C48/80 as measured by voltammetry. Two-way ANOVA showed a main effect of PBDE exposure (F (1, 22) = 6.3, *p* = 0.02). Fisher’s LSD revealed that the effect was driven by males, in which high PBDE compared to controls showed increased cumulative serotonin release (p=0.02). **(E)** β-hexosaminidase assay (n = 3/group). Two way ANOVA showed an interaction effect between source (pellet or supernatant) and Ionomycin stimulation (F (2, 12) = 4.2, *p* = 0.04). Fisher’s LSD revealed that pellets of high PBDE BMMCs showed reduced β-hexosaminidase content compared to both controls (p=0.001) and low PBDE (p=0.002).

**Supplementary figure 2.**
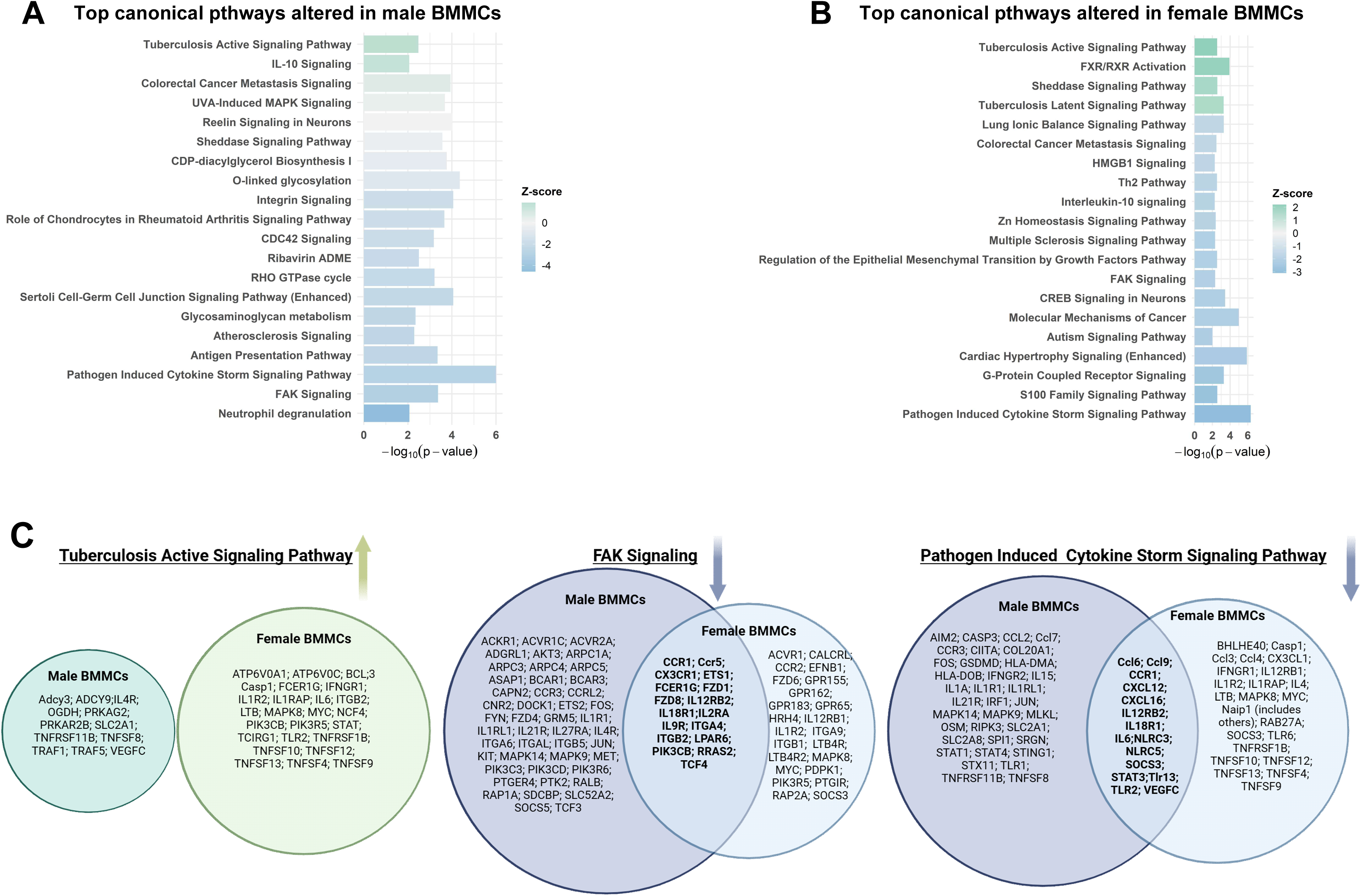
Ingenuity Pathway Analysis (IPA) of canonical pathways significantly enriched in male **(A)** and female **(B)** BMMCs. Shown are the top 20 pathways ranked by –log10(p-value). Bar colors reflect activation Z-score**.(C)** Selected upstream regulators predicted to be activated or inhibited in response to PBDE exposure in male and female BMMCs. Color indicates Z-score direction; bolded gene symbols represent regulators also differentially expressed at the transcript level in the corresponding sex.

## Notes

https://www.ncbi.xyz/geo/query/acc.cgi?acc=GSM8721796

### Competing Interest Statement

The authors have declared no competing interest.

### Summary of Updates

Link to GEO repository has been added, change in title

## References

1. Siddiqi, M. A., Laessig, R. H. & Reed, K. D. Polybrominated Diphenyl Ethers (PBDEs): New Pollutants-Old Diseases. Clinical Medicine and Research 1, 281 (2003).

2. Pollutants, P. O. Stockholm Convention on Persistent Organic Pollutants. (Geneva, 2011).

3. Sharkey, M., Harrad, S., Abou-Elwafa Abdallah, M., Drage, D. S. & Berresheim, H. Phasing-out of legacy brominated flame retardants: The UNEP Stockholm Convention and other legislative action worldwide. Environment International 144, 106041 (2020).

4. Saxena, P., Song, X., Zhang, B., Sarkar, A. & Achari, G. Profiling PBDE emissions from coastal landfills: Impact of waste management practices. Waste Management Bulletin 3, 391–401 (2025).

5. Zhang, Y., Xi, B. & Tan, W. Release, transformation, and risk factors of polybrominated diphenyl ethers from landfills to the surrounding environments: A review. Environment International 157, 106780 (2021).

6. Babichuk, N. et al. Polybrominated Diphenyl Ethers (PBDEs) in Marine Fish and Dietary Exposure in Newfoundland. EcoHealth 19, 99–113 (2022).

7. Turner, A. PBDEs in the marine environment: Sources, pathways and the role of microplastics. Environmental Pollution 301, 118943 (2022).

8. Bartalini, A., Muñoz-Arnanz, J., García-Álvarez, N., Fernández, A. & Jiménez, B. Global PBDE contamination in cetaceans. A critical review. Environ Pollut 308, 119670 (2022).

9. Sun, R.-X. et al. Evidence of polybrominated diphenyl ethers (PBDEs) and alternative halogenated flame retardants (AHFRs) in wild fish species from the remote tropical marine environment, south China sea. Environmental Pollution 361, 124885 (2024).

10. McGrath, T. J., Morrison, P. D., Sandiford, C. J., Ball, A. S. & Clarke, B. O. Widespread polybrominated diphenyl ether (PBDE) contamination of urban soils in Melbourne, Australia. Chemosphere 164, 225–232 (2016).

11. Lupton, S. J. Polybrominated diphenyl ethers (PBDEs) in US meat, poultry, and siluriformes: 2018–19 levels, trends, and estimated consumer exposures. Food Additives & Contaminants: Part A 42, 452–464 (2025).

12. Fernandes, A. R. et al. The transfer of environmental contaminants (Brominated and Chlorinated dioxins and biphenyls, PBDEs, HBCDDs, PCNs and PFAS) from recycled materials used for bedding to the eggs and tissues of chickens. Science of The Total Environment 892, 164441 (2023).

13. Olisah, C. et al. Extremely high levels of PBDEs in children’s toys from European markets: causes and implications for the circular economy. Environ Sci Eur 36, 1–11 (2024).

14. Li, Y. et al. Occurrence, levels and profiles of brominated flame retardants in daily-use consumer products on the Chinese market. Environ. Sci.: Processes Impacts 21, 446–455 (2019).

15. Kajiwara, N. et al. Recycling plastics containing decabromodiphenyl ether into new consumer products including children’s toys purchased in Japan and seventeen other countries. Chemosphere 289, 133179 (2022).

16. Liu, M., Brandsma, S. H. & Schreder, E. From e-waste to living space: Flame retardants contaminating household items add to concern about plastic recycling. Chemosphere 365, 143319 (2024).

17. Frederiksen, M., Vorkamp, K., Thomsen, M. & Knudsen, L. E. Human internal and external exposure to PBDEs--a review of levels and sources. Int J Hyg Environ Health 212, 109–134 (2009).

18. Klinčić, D., Dvoršćak, M., Jagić, K., Mendaš, G. & Herceg Romanić, S. Levels and distribution of polybrominated diphenyl ethers in humans and environmental compartments: a comprehensive review of the last five years of research. Environ Sci Pollut Res 27, 5744–5758 (2020).

19. Estill, C. F. et al. Biomarkers of Organophosphate and Polybrominated Diphenyl Ether (PBDE) Flame Retardants of American Workers and Associations with Inhalation and Dermal Exposures. Environ. Sci. Technol. 58, 8417–8431 (2024).

20. Hoffman, K. et al. Children’s exposure to brominated flame retardants in the home: The TESIE study. Environmental Pollution 352, 124110 (2024).

21. Zhang, Q. et al. Associations between polybrominated diphenyl ethers (PBDEs) levels in adipose tissues and blood lipids in women of Shantou, China. Environmental Research 214, 114096 (2022).

22. Giulivo, M. et al. Ecological and human exposure assessment to PBDEs in Adige River. Environmental Research 164, 229–240 (2018).

23. Hammel, S. C. et al. Novel and legacy brominated flame retardants in human breast milk and house dust from Denmark. Journal of Environmental Exposure Assessment 3, (2024).

24. Toms, L.-M. L. et al. Serum Polybrominated Diphenyl Ether (PBDE) Levels Are Higher in Children (2–5 Years of Age) than in Infants and Adults. Environmental Health Perspectives 117, 1461–1465 (2009).

25. Schecter, A. et al. Polybrominated Diphenyl Ether Flame Retardants in the U.S. Population: Current Levels, Temporal Trends, and Comparison With Dioxins, Dibenzofurans, and Polychlorinated Biphenyls. Journal of Occupational and Environmental Medicine 47, 199 (2005).

26. Toms, L.-M. L. et al. Higher accumulation of polybrominated diphenyl ethers in infants than in adults. Environ Sci Technol 42, 7510–7515 (2008).

27. Wallenborn, J. T. et al. Prenatal exposure to polybrominated diphenyl ether (PBDE) and child neurodevelopment: The role of breastfeeding duration. Science of The Total Environment 921, 171202 (2024).

28. Gaballah, S. et al. Distribution of polybrominated diphenyl ethers (PBDEs) in placental tissues of maternal and fetal origin in exposed Wistar rats and associations with thyroid hormone levels. Toxicological Sciences 204, 20–30 (2025).

29. Liu, Y. et al. Exposure levels and determinants of placental polybrominated diphenyl ethers in Chinese pregnant women. Environmental Research 241, 117615 (2024).

30. Ruis, M. T. et al. PBDEs Concentrate in the Fetal Portion of the Placenta: Implications for Thyroid Hormone Dysregulation. Endocrinology 160, 2748–2758 (2019).

31. Nanes, J. A. et al. Selected persistent organic pollutants in human placental tissue from the United States. Chemosphere 106, 20–27 (2014).

32. Zota, A. R. et al. Polybrominated diphenyl ethers (PBDEs) and hydroxylated PBDE metabolites (OH-PBDEs) in maternal and fetal tissues, and associations with fetal cytochrome P450 gene expression. Environment International 112, 269–278 (2018).

33. Stapleton, H. M., Kelly, S. M., Allen, J. G., McClean, M. D. & Webster, T. F. Measurement of Polybrominated Diphenyl Ethers on Hand Wipes: Estimating Exposure from Hand-to-Mouth Contact. Environ. Sci. Technol. 42, 3329–3334 (2008).

34. Kodavanti, P. R. S. et al. Developmental Exposure to a Commercial PBDE Mixture, DE-71: Neurobehavioral, Hormonal, and Reproductive Effects. Toxicological Sciences 116, 297–312 (2010).

35. Kuriyama, S. N., Talsness, C. E., Grote, K. & Chahoud, I. Developmental Exposure to Low-Dose PBDE-99: Effects on Male Fertility and Neurobehavior in Rat Offspring. Environmental Health Perspectives 113, 149–154 (2005).

36. Chen, A., Chung, E., DeFranco, E. A., Pinney, S. M. & Dietrich, K. N. Serum PBDEs and age at menarche in adolescent girls: analysis of the National Health and Nutrition Examination Survey 2003-2004. Environ Res 111, 831–837 (2011).

37. Harley, K. G. et al. Association of prenatal and childhood PBDE exposure with timing of puberty in boys and girls. Environment International 100, 132–138 (2017).

38. Lilienthal, H., Hack, A., Roth-Härer, A., Grande, S. W. & Talsness, C. E. Effects of Developmental Exposure to 2,2′,4,4′,5-Pentabromodiphenyl Ether (PBDE-99) on Sex Steroids, Sexual Development, and Sexually Dimorphic Behavior in Rats. Environmental Health Perspectives 114, 194–201 (2006).

39. Choi, G. et al. Polybrominated diphenyl ethers and incident pregnancy loss: The LIFE Study. Environ Res 168, 375–381 (2019).

40. Qiu, H. et al. Perinatal exposure to low-level PBDE-47 programs gut microbiota, host metabolism and neurobehavior in adult rats: An integrated analysis. Science of The Total Environment 825, 154150 (2022).

41. Strunz, S. et al. Maternal Exposure to Low-Dose BDE-47 Induced Weight Gain and Impaired Insulin Sensitivity in the Offspring. International Journal of Molecular Sciences 25, 8620 (2024).

42. Guo, J. et al. Umbilical cord serum PBDE concentrations and child adiposity measures at 7 years. Ecotoxicology and Environmental Safety 203, 111009 (2020).

43. Vuong, A. M. et al. Polybrominated diphenyl ether (PBDE) exposures and thyroid hormones in children at age 3Lyears. Environ Int 117, 339–347 (2018).

44. Lam, J. et al. Developmental PBDE Exposure and IQ/ADHD in Childhood: A Systematic Review and Meta-analysis. Environmental Health Perspectives 125, 086001 (2017).

45. Costa, L. G. & Giordano, G. DEVELOPMENTAL NEUROTOXICITY OF POLYBROMINATED DIPHENYL ETHER (PBDE) FLAME RETARDANTS. Neurotoxicology 28, 1047–1067 (2007).

46. Rice, D. C. et al. Developmental delays and locomotor activity in the C57BL6/J mouse following neonatal exposure to the fully-brominated PBDE, decabromodiphenyl ether. Neurotoxicology and Teratology 29, 511–520 (2007).

47. Bakhashab, S. et al. Acute and prolonged effects of interleukin-33 on cytokines in human cord blood-derived mast cells. Immunology Letters 269, 106908 (2024).

48. Conti, P. et al. Impact of TNF and IL-33 Cytokines on Mast Cells in Neuroinflammation. International Journal of Molecular Sciences 25, 3248 (2024).

49. Katsanos, G. S. et al. Mast cells and chemokines. J Biol Regul Homeost Agents 22, 145–151 (2008).

50. de Almeida, A. D. et al. A role for mast cells and mast cell tryptase in driving neutrophil recruitment in LPS-induced lung inflammation via protease-activated receptor 2 in mice. Inflamm Res 69, 1059–1070 (2020).

51. Mekori, Y. A. & Hershko, A. Y. T Cell-Mediated Modulation of Mast Cell Function: Heterotypic Adhesion-Induced Stimulatory or Inhibitory Effects. Front Immunol 3, 6 (2012).

52. Cao, J. et al. Human mast cells express corticotropin-releasing hormone (CRH) receptors and CRH leads to selective secretion of vascular endothelial growth factor. J Immunol 174, 7665–7675 (2005).

53. Ayyadurai, S. et al. Frontline Science: Corticotropin-releasing factor receptor subtype 1 is a critical modulator of mast cell degranulation and stress-induced pathophysiology. Journal of Leukocyte Biology 102, 1299–1312 (2017).

54. Narita, S. et al. Environmental Estrogens Induce Mast Cell Degranulation and Enhance IgE-Mediated Release of Allergic Mediators. Environ Health Perspect 115, 48–52 (2007).

55. McCallion, A. et al. Estrogen mediates inflammatory role of mast cells in endometriosis pathophysiology. Front. Immunol. 13, (2022).

56. Laoharatchatathanin, T. et al. Mast Cell Dynamics in the Ovary Are Governed by GnRH and Prolactin. Endocrinology 164, bqad144 (2023).

57. Alim, M. A. et al. Glutamate triggers the expression of functional ionotropic and metabotropic glutamate receptors in mast cells. Cell Mol Immunol 18, 2383–2392 (2021).

58. Suzuki, H., Miura, S., Liu, Y. Y., Tsuchiya, M. & Ishii, H. Substance P induces degranulation of mast cells and leukocyte adhesion to venular endothelium. Peptides 16, 1447–1452 (1995).

59. Xu, H. et al. Neurotransmitter and neuropeptide regulation of mast cell function: a systematic review. Journal of Neuroinflammation 17, 356 (2020).

60. Singh, T. S. K., Lee, S., Kim, H.-H., Choi, J. K. & Kim, S.-H. Perfluorooctanoic acid induces mast cell-mediated allergic inflammation by the release of histamine and inflammatory mediators. Toxicol Lett 210, 64– 70 (2012).

61. Miller, C. S., Palmer, R. F., Dempsey, T. T., Ashford, N. A. & Afrin, L. B. Mast cell activation may explain many cases of chemical intolerance. Environ Sci Eur 33, 129 (2021).

62. Park, S.-J., Sim, K. H., Shrestha, P., Yang, J.-H. & Lee, Y. J. Perfluorooctane sulfonate and bisphenol A induce a similar level of mast cell activation via a common signaling pathway, Fyn-Lyn-Syk activation. Food and Chemical Toxicology 156, 112478 (2021).

63. McCurdy, J. D., Lin, T. J. & Marshall, J. S. Toll-like receptor 4-mediated activation of murine mast cells. J Leukoc Biol 70, 977–984 (2001).

64. Mukai, K., Tsai, M., Saito, H. & Galli, S. J. Mast cells as sources of cytokines, chemokines, and growth factors. Immunol Rev 282, 121–150 (2018).

65. Shaik-Dasthagirisaheb, Y. B. et al. Vascular endothelial growth factor (VEGF), mast cells and inflammation. Int J Immunopathol Pharmacol 26, 327–335 (2013).

66. Yabut, J. M. et al. Genetic deletion of mast cell serotonin synthesis prevents the development of obesity and insulin resistance. Nat Commun 11, 463 (2020).

67. Borriello, F., Iannone, R. & Marone, G. Histamine Release from Mast Cells and Basophils. Handb Exp Pharmacol 241, 121–139 (2017).

68. Boyce, J. A. Mast cells and eicosanoid mediators: a system of reciprocal paracrine and autocrine regulation. Immunol Rev 217, 168–185 (2007).

69. Hochdörfer, T. et al. LPS-induced production of TNF-α and IL-6 in mast cells is dependent on p38 but independent of TTP. Cell Signal 25, 1339–1347 (2013).

70. Yuan, H. et al. Role of mast cell activation in inducing microglial cells to release neurotrophin. Journal of Neuroscience Research 88, 1348–1354 (2010).

71. Bao, C. et al. A mast cell–thermoregulatory neuron circuit axis regulates hypothermia in anaphylaxis. Science Immunology 8, eadc9417 (2023).

72. Heidarzadeh-Asl, S. et al. Novel insights on the biology and immunologic effects of histamine: A road map for allergists and mast cell biologists. Journal of Allergy and Clinical Immunology 155, 1095–1114 (2025).

73. Gao, S. et al. Histamine induced high mobility group box-1 release from vascular endothelial cells through H1 receptor. Front. Immunol. 13, (2022).

74. Barman, S. A. & Taylor, A. E. Histamine’s effect on pulmonary vascular resistance and compliance at elevated tone. American Journal of Physiology-Heart and Circulatory Physiology 257, H618–H625 (1989).

75. Okuno, T., Yabuki, A., Shiraishi, M., Obi, T. & Miyamoto, A. Histamine-induced modulation of vascular tone in the isolated chicken basilar artery: A possible involvement of endothelium. Comparative Biochemistry and Physiology Part C: Toxicology & Pharmacology 147, 339–344 (2008).

76. Dawodu, D., Sand, S., Nikolouli, E., Werfel, T. & Mommert, S. The mRNA expression and secretion of granzyme B are up-regulated via the histamine H2 receptor in human CD4+ T cells. Inflamm Res 72, 1525– 1538 (2023).

77. Takahashi, H. K. et al. Histamine downregulates CD14 expression via H2 receptorson human monocytes. Clinical Immunology 108, 274–281 (2003).

78. Takahashi, H. K. et al. Histamine inhibits lipopolysaccharide-induced interleukin (IL)-18 production in human monocytes. Clin Immunol 112, 30–34 (2004).

79. Hansson, M. et al. Histamine Protects T Cells and Natural Killer Cells Against Oxidative Stress. Journal of Interferon & Cytokine Research 19, 1135–1144 (1999).

80. Krishna, A., Beesley, K. & Terranova, P. F. Histamine, mast cells and ovarian function. J Endocrinol 120, 363–371 (1989).

81. Szukiewicz, D., Szewczyk, G., Klimkiewicz, J., Pyzlak, M. & Maslinska, D. The role of histamine and its receptors in the development of ovarian follicles in vitro. Inflamm. res. 55, S49–S50 (2006).

82. Sadek, A., Khramtsova, Y. & Yushkov, B. Mast Cells as a Component of Spermatogonial Stem Cells’ Microenvironment. Int J Mol Sci 25, 13177 (2024).

83. Frungieri, M. B. & Mayerhofer, A. Biogenic amines in the testis: sources, receptors and actions. Front. Endocrinol. 15, (2024).

84. Misto, A., Provensi, G., Vozella, V., Passani, M. B. & Piomelli, D. Mast Cell-Derived Histamine Regulates Liver Ketogenesis via Oleoylethanolamide Signaling. Cell Metabolism 29, 91–102.e5 (2019).

85. Ashina, K. et al. Histamine Induces Vascular Hyperpermeability by Increasing Blood Flow and Endothelial Barrier Disruption In Vivo. PLoS One 10, e0132367 (2015).

86. Atsushi, Y., Kazutaka, M., Takehiko, W., Hiroshi, W. & Yukihiko, K. Tissue distribution of histamine in a mutant mouse deficient in mast cells: Clear evidence for the presence of non-mast-cell histamine. Biochemical Pharmacology 31, 305–309 (1982).

87. Nautiyal, K. M., Ribeiro, A. C., Pfaff, D. W. & Silver, R. Brain mast cells link the immune system to anxiety-like behavior. Proc Natl Acad Sci U S A 105, 18053–18057 (2008).

88. Chikahisa, S. et al. Histamine from Brain Resident MAST Cells Promotes Wakefulness and Modulates Behavioral States. PLOS ONE 8, e78434 (2013).

89. Ramírez-Ponce, M. P. et al. Mast Cell Changes the Phenotype of Microglia via Histamine and ATP. Cell Physiol Biochem 55, 17–32 (2021).

90. Lenz, K. M. et al. Mast Cells in the Developing Brain Determine Adult Sexual Behavior. J Neurosci 38, 8044–8059 (2018).

91. Matsumoto, I., Inoue, Y., Shimada, T. & Aikawa, T. Brain Mast Cells Act as an Immune Gate to the Hypothalamic-Pituitary-Adrenal Axis in Dogs. J Exp Med 194, 71–78 (2001).

92. Bugajski, A. J., Chłap, Z., Gadek-Michalska, A., Borycz, J. & Bugajski, J. Degranulation and decrease in histamine levels of thalamic mast cells coincides with corticosterone secretion induced by compound 48/80. Inflamm Res 44 Suppl 1, S50–51 (1995).

93. Vizcaino, E., Grimalt, J. O., Fernández-Somoano, A. & Tardon, A. Transport of persistent organic pollutants across the human placenta. Environment International 65, 107–115 (2014).

94. Leonetti, C., Butt, C. M., Hoffman, K., Miranda, M. L. & Stapleton, H. M. Concentrations of Polybrominated Diphenyl Ethers (PBDEs) and 2,4,6-Tribromophenol in Human Placental Tissues. Environ Int 88, 23–29 (2016).

95. Kozlova, E. V. et al. Maternal transfer of environmentally relevant polybrominated diphenyl ethers (PBDEs) produces a diabetic phenotype and disrupts glucoregulatory hormones and hepatic endocannabinoids in adult mouse female offspring. Sci Rep 10, 18102 (2020).

96. Wang, D. et al. In utero and lactational exposure to BDE-47 promotes obesity development in mouse offspring fed a high-fat diet: impaired lipid metabolism and intestinal dysbiosis. Arch Toxicol 92, 1847–1860 (2018).

97. Woods, R. et al. Long-lived epigenetic interactions between perinatal PBDE exposure and Mecp2308 mutation. Hum Mol Genet 21, 2399–2411 (2012).

98. Seibenhener, M. L. & Wooten, M. C. Use of the Open Field Maze to Measure Locomotor and Anxiety-like Behavior in Mice. J Vis Exp 52434 (2015) doi:10.3791/52434.

99. Duque-Wilckens, N. et al. Oxytocin Receptors in the Anteromedial Bed Nucleus of the Stria Terminalis Promote Stress-Induced Social Avoidance in Female California Mice. Biol Psychiatry 83, 203–213 (2018).

100. Duque-Wilckens, N. et al. Extrahypothalamic oxytocin neurons drive stress-induced social vigilance and avoidance. Proc Natl Acad Sci U S A 117, 26406–26413 (2020).

101. Walf, A. A. & Frye, C. A. The use of the elevated plus maze as an assay of anxiety-related behavior in rodents. Nat Protoc 2, 322–328 (2007).

102. Mackey, E. et al. Perinatal androgens organize sex differences in mast cells and attenuate anaphylaxis severity into adulthood. Proceedings of the National Academy of Sciences 117, 23751–23761 (2020).

103. Duque-Wilckens, N. et al. Activity-dependent FosB gene expression negatively regulates mast cell functions. 2024.05.06.592755 Preprint at 10.1101/2024.05.06.592755 (2024).

104. D’Costa, S. et al. Mast cell corticotropin-releasing factor subtype 2 suppresses mast cell degranulation and limits the severity of anaphylaxis and stress-induced intestinal permeability. Journal of Allergy and Clinical Immunology 143, 1865–1877.e4 (2019).

105. Dobin, A. et al. STAR: ultrafast universal RNA-seq aligner. Bioinformatics 29, 15–21 (2013).

106. Anders, S., Pyl, P. T. & Huber, W. HTSeq--a Python framework to work with high-throughput sequencing data. Bioinformatics 31, 166–169 (2015).

107. Love, M. I., Huber, W. & Anders, S. Moderated estimation of fold change and dispersion for RNA-seq data with DESeq2. Genome Biology 15, 550 (2014).

108. Gülen, T. & Akin, C. Anaphylaxis and Mast Cell Disorders. Immunology and Allergy Clinics 42, 45–63 (2022).

109. Banafea, G. H., Bakhashab, S., Alshaibi, H. F., Natesan Pushparaj, P. & Rasool, M. The role of human mast cells in allergy and asthma. Bioengineered 13, 7049–7064 (2022).

110. Feyerabend, T. B. et al. Cre-mediated cell ablation contests mast cell contribution in models of antibody-and T cell-mediated autoimmunity. Immunity 35, 832–844 (2011).

111. Mackey, E. et al. Sexual dimorphism in the mast cell transcriptome and the pathophysiological responses to immunological and psychological stress. Biol Sex Differ 7, 60 (2016).

112. Duque-Wilckens, N. et al. Early life adversity drives sex-specific anhedonia and meningeal immune gene expression through mast cell activation. Brain Behav Immun 103, 73–84 (2022).

113. Roy, S., Na Ayudhya, C. C., Thapaliya, M., Deepak, V. & Ali, H. Multifaceted MRGPRX2: New Insight into the Role of Mast Cells in Health and Disease. J Allergy Clin Immunol 148, 293–308 (2021).

114. Mi, Y.-N., Ping, N.-N. & Cao, Y.-X. Ligands and Signaling of Mas-Related G Protein-Coupled Receptor-X2 in Mast Cell Activation. in Reviews of Physiology, Biochemistry and Pharmacology 139–188 (Springer, Cham, 2020). doi:10.1007/112_2020_53.

115. Ind, P. W., Brown, M. J., Lhoste, F. J. M., Macquin, I. & Dollery, C. T. Concentration effect relationships of infused histamine in normal volunteers. Agents and Actions 12, 12–16 (1982).

116. Brenner, B. et al. Plasma serotonin levels and the platelet serotonin transporter. J Neurochem 102, 206–215 (2007).

117. Dwyer, D. F., Barrett, N. A., Austen, K. F. & Consortium, T. I. G. P. Expression profiling of constitutive mast cells reveals a unique identity within the immune system. Nature immunology 17, 878 (2016).

118. Akula, S. et al. Quantitative In-Depth Analysis of the Mouse Mast Cell Transcriptome Reveals Organ-Specific Mast Cell Heterogeneity. Cells 9, 211 (2020).

119. Babina, M., Franke, K. & Bal, G. How “Neuronal” Are Human Skin Mast Cells? Int J Mol Sci 23, 10871 (2022).

120. Derakhshan, T., Boyce, J. A. & Dwyer, D. F. Defining mast cell differentiation and heterogeneity through single-cell transcriptomics analysis. Journal of Allergy and Clinical Immunology 150, 739–747 (2022).

121. Kitamura, Y., Shimada, M., Hatanaka, K. & Miyano, Y. Development of mast cells from grafted bone marrow cells in irradiated mice. Nature 268, 442–443 (1977).

122. Kitamura, Y., Go, S. & Hatanaka, K. Decrease of Mast Cells in W/Wv Mice and Their Increase by Bone Marrow Transplantation. Blood 52, 447–452 (1978).

123. Duque-Wilckens, N., et al. Mast cell-specific inactivation of Fosb exacerbates release of pro-inflammatory mediators in models of systemic anaphylaxis and lipopolysaccharide-induced sepsis. (2021).

124. Puebla-Osorio, N., Sarchio, S. N. E., Ullrich, S. E. & Byrne, S. N. Detection of Infiltrating Mast Cells Using a Modified Toluidine Blue Staining. Methods Mol Biol 1627, 213–222 (2017).

125. Ribatti, D. The Staining of Mast Cells: A Historical Overview. Int Arch Allergy Immunol 176, 55–60 (2018).

126. Fukuishi, N. et al. Does β-hexosaminidase function only as a degranulation indicator in mast cells? The primary role of β-hexosaminidase in mast cell granules. J Immunol 193, 1886–1894 (2014).

127. Cohen, R., Holowka, D. A. & Baird, B. A. Real-time Imaging of Ca2+ Mobilization and Degranulation in Mast Cells. Methods Mol Biol 1220, 347–363 (2015).

128. Hong-Tao, M. & Beaven, M. A. REGULATORS OF CA2+ SIGNALING IN MAST CELLS Potential Targets for Treatment of Mast-Cell Related Diseases? in Madame Curie Bioscience Database [Internet] (Landes Bioscience, 2013).

129. Cohen, R., Torres, A., Ma, H.-T., Holowka, D. & Baird, B. Ca2+ Waves Initiate Antigen-Stimulated Ca2+ Responses in Mast Cells. J Immunol 183, 6478–6488 (2009).

130. Omari, S., et al. Mast cell secretory granule fusion with amphisomes coordinates their homotypic fusion and release of exosomes. Cell Reports 43, (2024).

131. Sagi-Eisenberg, R. Biogenesis and homeostasis of mast cell lysosome related secretory granules. Front. Cell Dev. Biol. 13, (2025).

132. Aye, C. C., Toda, M., Morohoshi, K. & Ono, S. J. Identification of genes and proteins specifically regulated by costimulation of mast cell Fcε Receptor I and chemokine receptor 1. Exp Mol Pathol 92, 267–274 (2012).

133. Fifadara, N. H., Beer, F., Ono, S. & Ono, S. J. Interaction between activated chemokine receptor 1 and FcepsilonRI at membrane rafts promotes communication and F-actin-rich cytoneme extensions between mast cells. Int Immunol 22, 113–128 (2010).

134. Kuo, C.-H., Collins, A. M., Boettner, D. R., Yang, Y. & Ono, S. J. Role of CCL7 in Type I Hypersensitivity Reactions in Murine Experimental Allergic Conjunctivitis. J Immunol 198, 645–656 (2017).

135. Nagata, Y. & Suzuki, R. FcεRI: A Master Regulator of Mast Cell Functions. Cells 11, 622 (2022).

136. Turner, H. & Kinet, J.-P. Signalling through the high-affinity IgE receptor FcεRI. Nature 402, 24–30 (1999).

137. Brown, J., Jones, E. Y. & Forbes, B. E. Interactions of IGF-II with the IGF2R/cation-independent mannose-6-phosphate receptor mechanism and biological outcomes. Vitam Horm 80, 699–719 (2009).

138. Knight, P. A. et al. Aberrant Mucosal Mast Cell Protease Expression in the Enteric Epithelium of Nematode-Infected Mice Lacking the Integrin αvβ6, a Transforming Growth Factor-β1 Activator. The American Journal of Pathology 171, 1237–1248 (2007).

139. Welker, P., Grabbe, J., Gibbs, B., Zuberbier, T. & Henz, B. M. Nerve growth factor-β induces mast-cell marker expression during in vitro culture of human umbilical cord blood cells. Immunology 99, 418–426 (2000).

140. Aloe, L. The effect of nerve growth factor and its antibody on mast cells in vivo. Journal of Neuroimmunology 18, 1–12 (1988).

141. Baumann, C. A. & Saltiel, A. R. Spatial compartmentalization of signal transduction in insulin action. BioEssays 23, 215–222 (2001).

142. Halova, I., Draberova, L. & Draber, P. Mast Cell Chemotaxis – Chemoattractants and Signaling Pathways. Front. Immunol. 3, (2012).

143. Gilfillan, A. M. & Tkaczyk, C. Integrated signalling pathways for mast-cell activation. Nat Rev Immunol 6, 218–230 (2006).

144. Ferraro, F., Ma, X.-M., Sobota, J. A., Eipper, B. A. & Mains, R. E. Kalirin/Trio Rho Guanine Nucleotide Exchange Factors Regulate a Novel Step in Secretory Granule Maturation. MBoC 18, 4813–4825 (2007).

145. Taketomi, Y. & Murakami, M. Regulatory Roles of Phospholipase A2 Enzymes and Bioactive Lipids in Mast Cell Biology. Front. Immunol. 13, (2022).

146. Hong, H. et al. The Src family kinase Hck regulates mast cell activation by suppressing an inhibitory Src family kinase Lyn. Blood 110, 2511–2519 (2007).

147. Piliponsky, A. M. et al. Mast cell–derived factor XIIIA contributes to sexual dimorphic defense against group B streptococcal infections. J Clin Invest 132, (2022).

148. Etzel, R. A. The special vulnerability of children. International Journal of Hygiene and Environmental Health 227, 113516 (2020).

149. Yavropoulou, M. P., Sfikakis, P. P. & Chrousos, G. P. Immune System Effects on the Endocrine System. in Endotext (eds. Feingold, K. R. et al.) (MDText.com, Inc., South Dartmouth (MA), 2000).

150. Dai, M., Xu, Y., Gong, G. & Zhang, Y. Roles of immune microenvironment in the female reproductive maintenance and regulation: novel insights into the crosstalk of immune cells. Front. Immunol. 14, (2023).

151. Bilbo, S. D. & Schwarz, J. M. The Immune System and Developmental Programming of Brain and Behavior. Front Neuroendocrinol 33, 267–286 (2012).

152. Castellani, G., Croese, T., Peralta Ramos, J. M. & Schwartz, M. Transforming the understanding of brain immunity. Science 380, eabo7649 (2023).

153. Manley, K., Han, W., Zelin, G. & Lawrence, D. A. Crosstalk between the immune, endocrine, and nervous systems in immunotoxicology. Current Opinion in Toxicology 10, 37–45 (2018).

154. Chia, S. L., Kapoor, S., Carvalho, C., Bajénoff, M. & Gentek, R. Mast cell ontogeny: From fetal development to life-long health and disease. Immunol Rev 315, 31–53 (2023).

155. Baldwin, A. L. Mast cell activation by stress. Methods Mol Biol 315, 349–360 (2006).

156. Jiménez, M. et al. Responses of Mast Cells to Pathogens: Beneficial and Detrimental Roles. Front Immunol 12, 685865 (2021).

157. Gentek, R. et al. Hemogenic Endothelial Fate Mapping Reveals Dual Developmental Origin of Mast Cells. Immunity 48, 1160–1171.e5 (2018).

158. Dahlin, J. S. & Hallgren, J. Mast cell progenitors: Origin, development and migration to tissues. Molecular Immunology 63, 9–17 (2015).

159. Moon, T. C., Befus, A. D. & Kulka, M. Mast Cell Mediators: Their Differential Release and the Secretory Pathways Involved. Frontiers in Immunology 5, (2014).

160. Arowolo, O., Pilsner, J. R., Sergeyev, O. & Suvorov, A. Mechanisms of Male Reproductive Toxicity of Polybrominated Diphenyl Ethers. Int J Mol Sci 23, 14229 (2022).

161. Zhao, T. et al. Prenatal exposure to environmentally relevant levels of PBDE-99 leads to testicular dysgenesis with steroidogenesis disorders. Journal of Hazardous Materials 424, 127547 (2022).

162. Talsness, C. E. et al. In Utero and Lactational Exposures to Low Doses of Polybrominated Diphenyl Ether-47 Alter the Reproductive System and Thyroid Gland of Female Rat Offspring. Environ Health Perspect 116, 308–314 (2008).

163. Varshavsky, J. R. et al. Association of polybrominated diphenyl ether (PBDE) levels with biomarkers of placental development and disease during mid-gestation. Environmental Health 19, 61 (2020).

164. Harley, K. G. et al. PBDE Concentrations in Women’s Serum and Fecundability. Environmental Health Perspectives 118, 699–704 (2010).

165. Mondillo, C., Patrignani, Z., Reche, C., Rivera, E. & Pignataro, O. Dual role of histamine in modulation of Leydig cell steroidogenesis via HRH1 and HRH2 receptor subtypes. Biol Reprod 73, 899–907 (2005).

166. Pagotto, R. M., Monzón, C., Moreno, M. B., Pignataro, O. P. & Mondillo, C. Proliferative effect of histamine on MA-10 Leydig tumor cells mediated through HRH2 activation, transient elevation in cAMP production, and increased extracellular signal-regulated kinase phosphorylation levels. Biol Reprod 87, 150 (2012).

167. Mondillo, C., Varela, M. L., Abiuso, A. M. B. & Vázquez, R. Potential negative effects of anti-histamines on male reproductive function. (2018) doi:10.1530/REP-17-0685.

168. Woidacki, K., Jensen, F. & Zenclussen, A. C. Mast cells as novel mediators of reproductive processes. Front. Immunol. 4, (2013).

169. Woidacki, K. et al. Mast cells rescue implantation defects caused by c-kit deficiency. Cell Death Dis 4, e462 (2013).

170. Park, S. et al. Prenatal exposure to polybrominated diphenyl ethers and inattention/hyperactivity symptoms in mid to late adolescents. Front. Epidemiol. 3, (2023).

171. Chen, A. et al. Prenatal Polybrominated Diphenyl Ether Exposures and Neurodevelopment in U.S. Children through 5 Years of Age: The HOME Study. Environmental Health Perspectives 122, 856–862 (2014).

172. Vuong, A. M. et al. Childhood polybrominated diphenyl ether (PBDE) exposure and neurobehavior in children at 8 years. Environmental Research 158, 677–684 (2017).

173. Hartley, K. et al. Gestational exposure to polybrominated diphenyl ethers and social skills and problem behaviors in adolescents: The HOME study. Environment International 159, 107036 (2022).

174. Carthy, E. & Ellender, T. Histamine, Neuroinflammation and Neurodevelopment: A Review. Front. Neurosci. 15, (2021).

175. Mochizuki, T. Histamine as an Alert Signal in the Brain. in The Functional Roles of Histamine Receptors 413–425 (Springer, Cham, 2021). doi:10.1007/7854_2021_249.

176. Qian, H., Shu, C., Xiao, L. & Wang, G. Histamine and histamine receptors: Roles in major depressive disorder. Front. Psychiatry 13, (2022).

177. Rani, B. et al. Brain histamine and oleoylethanolamide restore behavioral deficits induced by chronic social defeat stress in mice. Neurobiol Stress 14, 100317 (2021).

178. Dropp, J. J. Mast cells in mammalian brain. Acta Anat (Basel) 94, 1–21 (1976).

179. Matsumoto, I., Inoue, Y., Shimada, T. & Aikawa, T. Brain mast cells act as an immune gate to the hypothalamic-pituitary-adrenal axis in dogs. J Exp Med 194, 71–78 (2001).

180. Silver, R., Silverman, A.-J., Vitković, L. & Lederhendler, I. I. Mast cells in the brain: evidence and functional significance. Trends in Neurosciences 19, 25–31 (1996).

181. Schlicker, E. & Kathmann, M. Role of the Histamine H3 Receptor in the Central Nervous System. Handb Exp Pharmacol 241, 277–299 (2017).

182. Kolkhir, P., Elieh-Ali-Komi, D., Metz, M., Siebenhaar, F. & Maurer, M. Understanding human mast cells: lesson from therapies for allergic and non-allergic diseases. Nat Rev Immunol 22, 294–308 (2022).

183. Akin, C. Mast cell activation syndromes. Journal of Allergy and Clinical Immunology 140, 349–355 (2017).

184. Hamilton, M. J., Frei, S. M. & Stevens, R. L. The Multifaceted Mast Cell in Inflammatory Bowel Disease. Inflamm Bowel Dis 20, 2364–2378 (2014).

185. McClain, J. L. et al. Histamine-dependent interactions between mast cells, glia, and neurons are altered following early-life adversity in mice and humans. Am J Physiol Gastrointest Liver Physiol 319, G655–G668 (2020).

186. Guan, L. C., Dong, X. & Green, D. P. Roles of mast cells and their interactions with the trigeminal nerve in migraine headache. Mol Pain 19, 17448069231181358 (2023).

187. Nurkhametova, D. F. et al. Mast Cell Mediators as Pain Triggers in Migraine: Comparison of Histamine and Serotonin in the Activation of Primary Afferents in the Meninges in Rats. Neurosci Behav Physi 50, 900– 906 (2020).

188. Hendriksen, E., van Bergeijk, D., Oosting, R. S. & Redegeld, F. A. Mast cells in neuroinflammation and brain disorders. Neurosci Biobehav Rev 79, 119–133 (2017).

189. Paul, W. E. & Zhu, J. How are TH2-type immune responses initiated and amplified? Nat Rev Immunol 10, 225–235 (2010).

190. Ryan, R. Y. M. et al. Immunological Responses to Envenomation. Front. Immunol. 12, (2021).

191. Zaiss, D. M. W., Pearce, E. J., Artis, D., McKenzie, A. N. J. & Klose, C. S. N. Cooperation of ILC2s and TH2 cells in the expulsion of intestinal helminth parasites. Nat Rev Immunol 24, 294–302 (2024).

192. Kabat, A. M. et al. Resident TH2 cells orchestrate adipose tissue remodeling at a site adjacent to infection. Science Immunology 7, eadd3263 (2022).

193. Goetzl, E. J. Th2 cells in rapid immune responses and protective avoidance reactions. The FASEB Journal 38, e23485 (2024).

194. Florsheim, E. B. et al. Immune sensing of food allergens promotes avoidance behaviour. Nature 620, 643–650 (2023).

195. Mazzoni, A., Siraganian, R. P., Leifer, C. A. & Segal, D. M. Dendritic Cell Modulation by Mast Cells Controls the Th1/Th2 Balance in Responding T Cells1. The Journal of Immunology 177, 3577–3581 (2006).

196. Dirscherl, C. et al. Dissociation of β2m from MHC class I triggers formation of noncovalent transient heavy chain dimers. Journal of Cell Science 135, jcs259489 (2022).

197. McShan, A. C. et al. TAPBPR promotes antigen loading on MHC-I molecules using a peptide trap. Nat Commun 12, 3174 (2021).

198. Garcia-Lora, A., Algarra, I. & Garrido, F. MHC class I antigens, immune surveillance, and tumor immune escape. Journal of Cellular Physiology 195, 346–355 (2003).

199. León Machado, J. A. & Steimle, V. The MHC Class II Transactivator CIITA: Not (Quite) the Odd-One-Out Anymore among NLR Proteins. International Journal of Molecular Sciences 22, 1074 (2021).

200. Douillard, V. et al. Approaching Genetics Through the MHC Lens: Tools and Methods for HLA Research. Front. Genet. 12, (2021).

201. Lotfi-Emran, S. et al. Human mast cells present antigen to autologous CD4+ T cells. Journal of Allergy and Clinical Immunology 141, 311–321.e10 (2018).

202. Stelekati, E. et al. Mast cell-mediated antigen presentation regulates CD8+ T cell effector functions. Immunity 31, 665–676 (2009).

203. Kambayashi, T. & Laufer, T. M. Atypical MHC class II-expressing antigen-presenting cells: can anything replace a dendritic cell? Nat Rev Immunol 14, 719–730 (2014).

204. Galli, S. J. & Gaudenzio, N. Human mast cells as antigen-presenting cells: When is this role important in vivo? Journal of Allergy and Clinical Immunology 141, 92–93 (2018).

205. Akintunde, M. E. et al. Ex vivo exposure to polybrominated diphenyl ether (PBDE) selectively affects the immune response in autistic children. *Brain, Behavior*, & Immunity - Health 34, 100697 (2023).

206. Hong, S. K. et al. Polybrominated Diphenyl Ethers Orally Administration to Mice Were Tansferred to Offspring during Gestation and Lactation with Disruptions on the Immune System. Immune Netw 10, 64–74 (2010).

207. Frouin, H., Lebeuf, M., Hammill, M., Masson, S. & Fournier, M. Effects of individual polybrominated diphenyl ether (PBDE) congeners on harbour seal immune cells *in vitro*. Marine Pollution Bulletin 60, 291–298 (2010).

